# New Structural Insights into the Function of the Catalytically Active Human Taspase1

**DOI:** 10.1101/2020.11.26.400622

**Authors:** Nirupa Nagaratnam, Silvia L. Delker, Rebecca Jernigan, Thomas E. Edwards, Janey Snider, Darren Thifault, Dewight Williams, Brent L. Nannenga, Mary Stofega, Lidia Sambucetti, James J. Hsieh, Andrew J. Flint, Petra Fromme, Jose M. Martin-Garcia

## Abstract

Proteases can play essential roles in severe human pathology, ranging from degenerative and inflammatory illnesses to infectious diseases, with some, such as Taspase1, involved in growth and progression of tumors at primary and metastatic sites. Taspase1 is a N-terminal nucleophile (Ntn)-hydrolase overexpressed in primary human cancers, coordinating cancer cell proliferation, invasion, and metastasis. Loss of Taspase1 activity disrupts proliferation of human cancer cells *in vitro* and in mouse xenograft models of glioblastoma, thus this protein has the potential to become a novel anticancer drug target. It belongs to the family of Ntn-hydrolases, a unique family of proteins synthesized as enzymatically inactive proenzymes that become activated upon cleavage of the peptide bond on the N-terminal side of a threonine residue, which then becomes the catalytic site nucleophile. The activation process simultaneously changes the conformation of a long domain at the C-terminus of the alpha-subunit for which no full-length structural information exists and its function is poorly understood. Here we present a novel cloning strategy to generate a fully active, circularly permuted form of Taspase1 to determine the crystallographic structure of catalytically active human Taspase1 to 3.04Å. We discovered that this region forms a long helical domain and is indispensable for the catalytic activity of Taspase1. Together, our study highlights the importance of this element for the enzymatic activity of Ntn-hydrolases and suggests that this long domain could be a novel target for the design of inhibitors with the potential to be developed into anticancer therapeutics.

## Introduction

Proteases are key enzymes in the human genome that carry out important biological processes including, but not limited to, metabolism, tissue remodeling, apoptosis, cell proliferation, and migration. Further, this class of enzymes is implicated in the progression and growth of tumors when their function is deregulated (1, 2). Taspase1 is an endopeptidase that promotes cell proliferation and permits oncogenic initiation through cleavage of the human mixed-lineage leukemia (MLL) nuclear protein. The MLL activation by Taspase1 has been shown to be a key step in several cancer types, including development of childhood leukemia (3, 4). Taspase1 is also involved in proper regulation of expression of *HOX* genes and regulation of cell cycle genes (5, 6). MLL is a large nuclear protein of 3,969 residues (500 kDa) that is synthesized in an inactive form. Hsieh and co-workers discovered that Taspase1 catalyzes the processing of MLL from its inactive to active state of the protein (4). Subsequent studies identified additional Taspase1 substrates, including MLL2, TFIIA, ALF (TFIIA), and Drosophila HCF (6-8). Taspase1 mediates cleavage of its substrates by recognizing conserved peptide motifs (Q^3^ X^2^ D^1^/G^-1^ X^-2^ D^-3^ D^-4^) with Asp and Gly residues at positions P1 and P-1, respectively (4). In the case of MLL, Taspase1 cleaves the protein at two sites (after D2666 and D2718), creating an N-terminal fragment of 320 kDa and a C-terminal fragment of 180 kDa, which heterodimerize to form a stable complex that localizes to a subnuclear compartment (4). Despite intensive work on Taspase1, the catalytic mechanism by which Taspase1 cleaves and thereby activates its very large target proteins, and the regulation of that activity leading to severe changes in gene expression that promote cancer development are still not well understood (9).

Taspase1 is a unique threonine aspartase that contains an N-terminal threonine residue as a nucleophile and cleaves after an aspartate residue in the target substrate (3, 4). The human *taspase1* gene encodes a highly conserved protein of 420 residues with a molecular weight of 50 kDa. Taspase1 belongs to a superfamily of proteases called N-terminal nucleophile hydrolases (Ntn hydrolases) (10) that are synthesized as enzymatically inactive proenzymes that become activated upon cleavage of the peptide bond on the N-terminal side of the active site threonine, serine or cysteine. In Taspase1, the intramolecular reaction takes place between residues Asp233 and Thr234 to generate an N-terminal fragment of 28 kDa (α subunit) and a C-terminal fragment of 22 kDa (β subunit). The N-terminal threonine residue of the β-subunit becomes the active site nucleophile(3) essential for proteolytic cleavage of substrates. The two subunits remain associated and assemble into a higher order αββα fold (10-13).

No structural information is available for Taspase1 in a complex with any of its target substrates. Only high-resolution crystal structures of the proenzyme and a mature, 2-chain truncated version of human Taspase1 have been determined so far (14). Due to the high flexibility of the region between residues Pro183 and Asp233, Khan *et al*. removed residues 207-233 to enable crystallization (14). Both the proenzyme and the truncated Taspase1 were demonstrated to be dimers, and the overall structure showed no major differences between the proenzyme dimer and the dimer of the truncated protein, with only some reorganization at the dimer interface and significant conformational differences of the side chains in or near the catalytic site, including the catalytic nucleophile Thr234, also were reported (14). Importantly, the cleavage to activate Taspase1 releases and increases the flexibility a 56 amino acid segment at the C-terminus of the alpha-subunit previously constrained by attachment to Thr234. But, in this conformation the sequence had only been partially resolved in prior crystal structures of Taspase1 (14). Interestingly, this loop is also absent in all crystal structures of type 2 asparaginases solved so far (10-13, 15-17).

In a recent study by van den Boom and co-workers, they used molecular modeling, CD, and NMR spectroscopy to predict that this long loop was mostly alpha-helical with a defined helix-turn-helix secondary structure (18). They also hypothesized that, in the context of the full length Taspase1, the long loop might be in a closed conformation, sitting on top of the catalytic site of the proenzyme, and then undergoing a large conformational change to adopt a more flexible status, or open state, in the active Taspase1 (18). The proximity of this unresolved region to the active site and its crucial role in the activation of the cancer-related protease Taspase1 suggest this region could be a novel target for Taspase1 inhibitors. One approach could be the design of inhibitors that bind the loop region acting as clamps to keep the loop in a closed conformation, preventing activation of the Taspase1 proenzyme.

Although published studies (14, 18) have provided important clues to the biological function of Taspase1, structural information on the active form of the enzyme will be critical to help us understand how Taspase1 recognizes and processes its natural targets. Here, we present a novel cloning strategy to generate a fully active, circularly permuted form of Taspase1 by swapping the α and β subunits of the protein so that the region with the long domain is placed at the C-terminus of a single-chain enzyme. This strategy has allowed us to determine the crystal structure of catalytically active human Taspase1 at 3.04Å resolution. The structure reveals the conformation of a clearly defined, long helical domain (Pro183-Asp233) in the open state of the active enzyme. In addition, we demonstrate that this long domain is essential for Taspase1 activity. Taken together, these results lead us to hypothesize that the current view of the proteolytic mechanism of Ntn-hydrolases must be revisited and the long domain has to be included to fully understand how this family of proteases recognize and process their targets.

## Results

### The circularly permuted constructs of human Taspase1

As Taspase1 undergoes post-translational autoproteolytic cleavage between residues Asp233 and Thr234 and forms a hetero-tetramer composed of alpha and beta subunits (3), isolation of recombinant Taspase1 in homogenous form is complicated by the presence of two species; the inactive proenzyme and the active cleaved enzyme. Because these two species have essentially the same molecular weight, separation by biophysical techniques is difficult, and obtaining pure protein for structural studies is challenging (14). In order to obtain homogenous Taspase1, Khan and co-workers created a mutant of the inactive proenzyme (D233A) and successfully purified and crystallized it. The crystal structure revealed two disordered regions in the alpha subunit, 1-40 and ∼203-233. To obtain Taspase1 in its processed cleaved form, they co-expressed and purified the associated alpha and beta subunits. However, attempts to crystallize the hetero-tetramer were unsuccessful due to the presence of disordered regions. A similar co-expression strategy in which residues 207-233 of the alpha subunit were deleted provided the first crystal structure of a mature, 2-chain truncated version of Taspase1 in its hetero-tetrameric form (14). However, residues 183-206 remained unresolved in this structure.

To obtain the first structure of human Taspase1 in its catalytically active state, we designed, expressed and purified three circularly permuted (cp) Taspase1 variants (cp-Taspase1*R*_α41-183/β_,cp-Taspase1_α41-206/β_,cp-Taspase1_α41-233/β_). These constructs have α and β subunits swapped and linked through a GSGS tetrapeptide linker. A comparison of the sequences of our cp constructs with the sequence of WT Taspase1 is shown in Figure 1A. In our constructs, the nucleophile Thr234 becomes the first residue of the N-terminus as it would be found in the wild type (WT) Taspase1 after autoproteolytic cleavage. We deleted the first 40 residues of the alpha subunit, which are disordered in previous structures (14), and linked this new N-terminus to the C-terminus of the β subunit using the GSGS linker described above. The region 183-233 is also shown to be disordered in the structure of Khan *et al*. However, this region lies adjacent to the catalytic site, the deletion of which might perturb the active site of the enzyme (14). Yet, minimizing disordered regions increases the probability of crystallization and structure determination. Thus, we performed serial truncations in the long domain of the alpha subunit (residues 183-233) in the cp constructs to study the effect of this domain on the catalytic function of the enzyme. Figure 1B shows a comparison of the cp constructs used in our study to the constructs previously reported (14) and to the WT 2-chain Taspase1.

**Figure 1.**
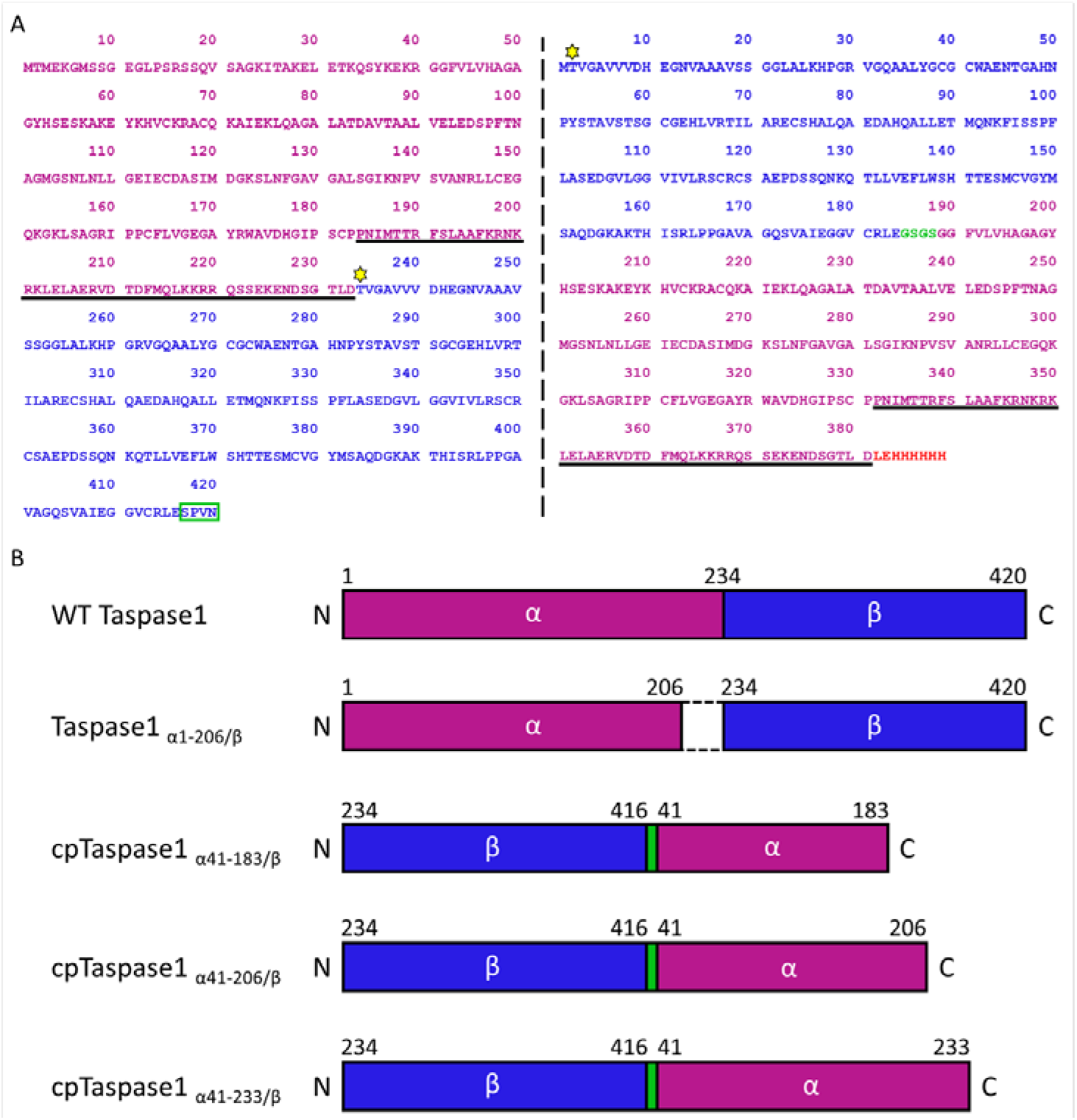
Sequence and domain organization of circularly permuted constructs. A) Comparison of the sequence of the circularly permuted construct, cpTaspase1_α41-233/β_ (right), with the sequence of the WT Taspase1 (left). The catalytic site nucleophile Thr234 is highlighted by the yellow star. The α and β subunits are colored in magenta and blue, respectively. In the circularly permuted construct, α and β subunits are swapped and linked through a GSGS linker (green). The GSGS residues replace the SPVN residues in the WT Taspase1 (green, boxed). The residues comprising the long C-terminal helix fragment are underlined. The first 40 residues of the alpha subunit are deleted except for the N-terminal initiating methionine. The two extra LE residues and the His (H) tag residues at the C-terminus (highlighted in red) are for cloning and for affinity purification purposes, respectively. B) Schematic representation of Taspase1 constructs. Amino acid positions in all constructs are numbered according to the wild type Taspase1 (WT Taspase1). The α and β subunits are highlighted in magenta and blue, respectively. Taspase1_α1-206/β_ construct (PDB 2A8J(14)) shows the two subunits are co-expressed with the missing fragment highlighted by the dashed lines. The three circularly permuted constructs (cp-Taspase1_α41-183/β_, cp-Taspase1_α41-206/β_, cp-Taspase1_α41-233/β_) used in our study are shown: both subunits are swapped and linked through a GSGS tetrapeptide (green). These circularly permuted constructs differ from each other only in the length of the alpha subunit sequence.

### Catalytic activity of the cp human Taspase1 proteins

To assess the catalytic activity of the cp Taspase1 proteins, we developed a quenched FRET-based assay that measures the proteolysis of a labeled Taspase1 peptide substrate. The internally quenched peptide substrate includes 10 residues (P_-6_ – P_4_) from cleavage site 2 (after D2718) in MLL, which is more efficiently processed by Taspase1 than cleavage site 1 (after D2666)(3). For the fluorescence emitter, we used an MCA (7-Methoxycoumarin-4-yl)acetyl) fluorophore attached to the N-terminus of the peptide through the first Lys residue; the quencher used was 2,4-dinitrophenyl-lysine (Lys(DNP)) attached to the lysine residue at the C terminus, which is not part of the native sequence in MLL. The peptide was also amidated at the N-terminus. The sequence of the peptide substrate was MCA-Lys-Ile-Ser-Gln-Leu-Asp↓Gly-Val-Asp-Asp-Lys(DNP)-NH_2_ where the cleavage site is indicated by the arrow (↓). The proteolytic cleavage of the substrate by Taspase1 proteins was monitored for ∼1.5 h after the addition of substrate. Figure 2 shows the reaction progress curves for the three cp proteins compared to that of the WT Taspase1. The assay without enzyme (substrate alone) was included as a negative control. We observed high catalytic activity for the WT Taspase1 and comparable or slightly higher activity for the long circular permuted construct cp-Taspase1_α41-233/β_. However, the catalytic activity of Taspase1 was completely abolished by deleting the fragment 183-233 in the shortest construct (cp-Taspase1_α41-183/β_ construct), giving a signal similar to the substrate alone. An intermediate catalytic activity was measured for the cp-Taspase1_α41-206/β_ protein. These results revealed that the region between 183 and 233 is essential for the catalytic activity of Taspase1, whereas previously it was assumed that the deletion of the fragment 183-233 would lead to a fully active enzyme (14). Thus, structural characterization of this functionally important fragment of the enzyme by crystallography became an even more important quest, as it might reveal features of the catalytic activity of Taspase1 that had previously been unappreciated.

**Figure 2.**
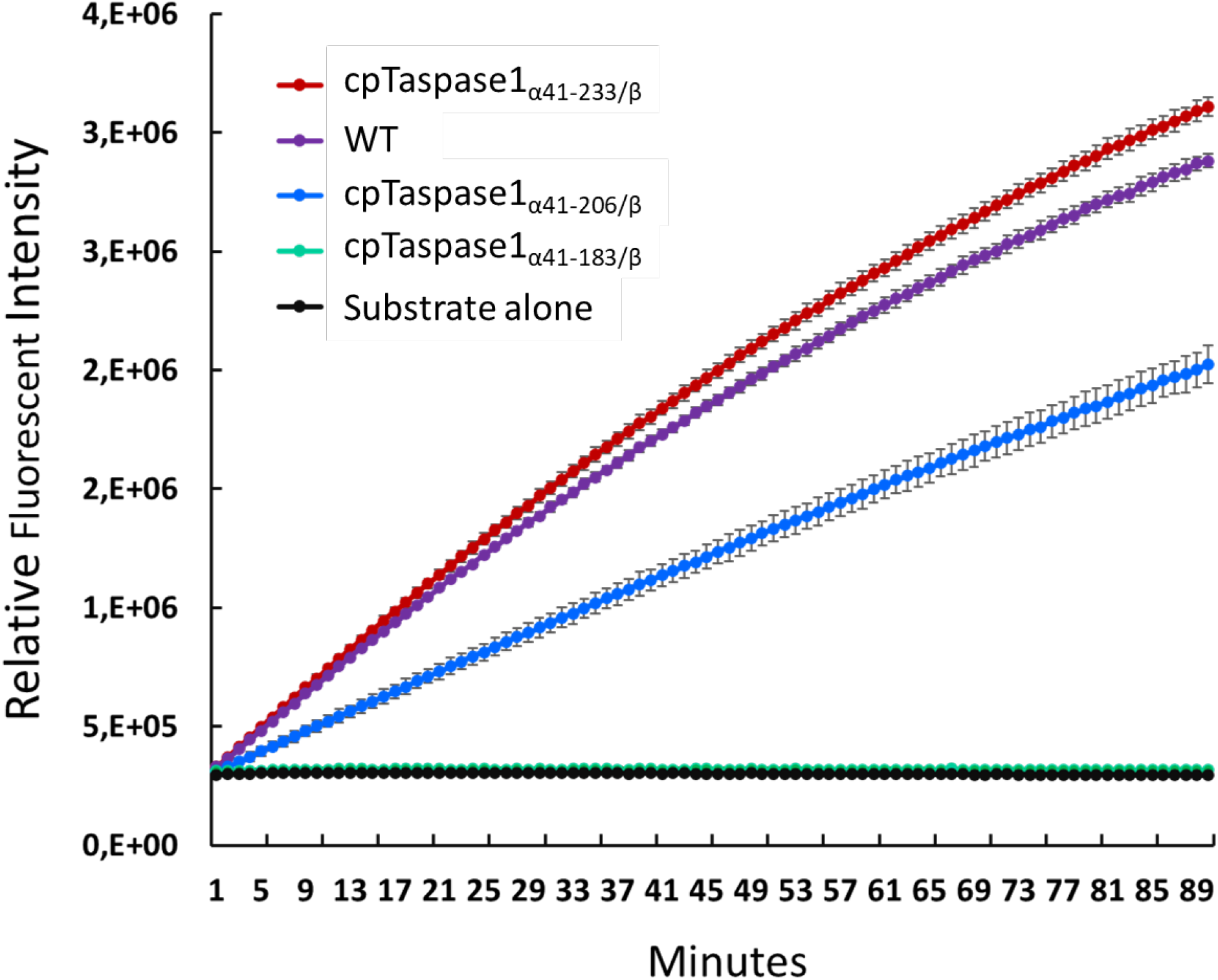
Activity assays of the circularly permuted constructs (cpTaspase1_α41-233/β_ (red), cpTaspase1_α41-206/β_ (blue), and cpTaspase1_α41-183/β_ (green)), and the WT Taspase1 (purple). Reaction progress curves of the proteolytic cleavage of the labeled peptidic substrate MCA-Lys-Ile-Ser-Gln-Leu-Asp↓Gly-Val-Asp-Asp-Lys-(DNP)-NH_2_ by various Taspase1 proteins. The experiment with the substrate alone (control) is represented in black. Error bars have been included.

### Structure determination of the short cp human Taspase1

To determine if the circular permutations induced any structural alteration compared to the structure of the mature, 2-chain truncated Taspase1 from Khan *et al*. (14), we solved the crystal structure of the shortest circularly permuted (cp) constructs (cp-Taspase1 _α41-183/β_). The medium length circularly permuted cp-Taspase1_α41-206/β_ construct was also purified, but crystallization experiments were unsuccessful, likely due to the high flexibility of the extra residues 183-206.

The crystal structure of the cp-Taspase1_α41-183/β_ protein was determined at 2.15 Å resolution (Table 1) by molecular replacement using the structure of the mature, 2-chain truncated Taspase1 reported by Khan and co-workers (PDB 2A8J) (14). The crystals of cp-Taspase1_α41-183/β_ belonged to the space group P6_5_. The crystals contain the characteristic hetero-tetramer in the asymmetric unit, as previously described for the mature, 2-chain truncated enzyme (14) (Figure S1A).

**Table 1.** Crystallographic data collection and refinement statistics.

|  | cp-Taspase I<sub>α41-233/β</sub> | cp-Taspase I<sub>α41-183/β</sub> |
| --- | --- | --- |
| ***Data collection statistics*** |  |  |
| Radiation source | APS 23-ID-D | APS 21-ID-F |
| Wavelength (Å) | 1.04 | 0.97 |
| Detector | PILATUS3 6M | RAYONIX<br>MX300HE |
| Temperature (K) | 100 | 100 |
| Space group | H32 | P6<sub>5</sub> |
| Unit cell dimensions |  |  |
| a, b, c (Å) | 196.00, 196.00, 196.91 | 60.30, 60.30, 319.01 |
| α, β, γ (°) | 90, 90, 120 | 90, 90, 120 |
| Resolution range (Å) | 3.04-49.05 (3.04-3.11) | 2.15-40.41 (2.15-2.20) |
| I/δ(I) | 17.0 (1.9) | 15.24 (3.74) |
| Multiplicity | 19.6 (20.4) | 5.8 (5.8) |
| Completeness (%) | 99.8 (99.4) | 99.8 (85.8) |
| CC<sub>1/2</sub> (%) | 100 (91.1) | 99.8 (85.8) |
| R<sub>merge</sub> (%) | 6.8 (187.5) | 7.8 (52.0) |
| Total number reflections | 343,892 | 206,406 |
| No. of unique reflections | 17,964 | 35,513 |
| ***Ellipsoidal truncation*** |  |  |
| Ellipsoidal<sup>a</sup> resolution (Å) (direction)<sup>b</sup> | 2.9 (0.894a* – 0.447b*)<br>2.9 (b*)<br>4.8 (c*) | N/A |
| Best/Worst diffraction limits after cut-off (Å) | 3.04 / 5.3 | N/A |
| I/δ(I) (ellipsoidal<sup>a</sup>)<sup>c</sup> | 21.2 (1.5) | N/A |
| Completeness (%) (spherical<sup>a</sup>)<sup>c,d</sup> | 54.4 (13.7) | N/A |
| Completeness (%) (ellipsoidal<sup>a</sup>)<sup>c,d</sup> | 93.9 (65.5) | N/A |
| Total no. of reflections (ellipsoidal<sup>a</sup>)<sup>c</sup> | 299,919 | N/A |
| No. of unique reflections (ellipsoidal<sup>a</sup>)<sup>c</sup> | 15,315 | N/A |
| ***Data refinement statistics for truncated data*** |  |  |
| Number of reflections used in refinement | 13,749 | 35,417 |
| Number of free reflections used in refinement | 1,568 | 1,709 |
| R<sub>work</sub> / R<sub>free</sub> (%) | 23.7 / 30.7 | 15.9 / 19.7 |
| Wilson B-factor (Å<sup>2</sup>) | 152.0 | 43.5 |
| Solvent content (%) / Matthews coefficient (V<sub>m</sub>) | 70 / 4.3 | 48 / 2.4 |
| No. of atoms |  |  |
| Protein | 10,510 | 4,368 |
| Water and others (ligands or ions) | 0 | 189 |
| Overall B factor (Å<sup>2</sup>) | 165.0 | 43.48 |
| R.M.S. deviations from ideal values |  |  |
| Bonds lengths (Å) | 0.007 | 0.007 |
| Bond angles (°) | 1.744 | 0.827 |
| Ramachandran plot statistics (%) |  |  |
| Favored | 85 | 96.55 |
| Allowed | 12 | 2.80 |
| Outliers | 3 | 0 |
| Molprobity score / percentile | 3.5 / 38<sup>th</sup> | 1.55 / 99<sup>th</sup> |
| PDB | 6VIN | 6UGK |

<sup>a</sup>These statistics are for data that were truncated by STARANISO to remove poorly measured reflections affected by anisotropy.
<sup>b</sup>The resolution limits for three directions in reciprocal space are indicated here. To accomplish this, STARANISO computed an ellipsoid postfitted by least squares to the cutoff surface, removing points where the fit was poor. Note that the cutoff surface is unlikely to be perfectly ellipsoidal, so this is only an estimate.
<sup>c</sup>Values in parentheses are for the highest-resolution shell. Note that data collection and refinement statistics have different highest-resolution shells. <sup>d</sup>The anisotropic completeness was obtained by least squares fitting an ellipsoid to the reciprocal lattice points at the cutoff surface defined by a local mean $I/\sigma I$ threshold of 1.2, rejecting outliers in the fit due to spurious deviations (including any cusp), and calculating the fraction of observed data lying inside the ellipsoid so defined. Note that the cutoff surface is unlikely to be perfectly ellipsoidal, so this is only an estimate.

The structural alignment of cp-Taspase1_α41-183/β_ with the mature, 2-chain truncated Taspase1 protein from Khan *et al*., showed that the overall structure of both inactive forms of Taspase1 are very similar to one another with average RMSDs of 0.44 Å and 0.60 Å for the C_α_ and all visible residues, respectively (Table S1). Thus, reassuringly, the circular permutation has not altered the general folding of the protein (Figure S1B). There are, however, minor differences in the loop regions between the two structures (Figure S1B). A comparison of the two chains in the asymmetric unit showed that the α and β subunits have essentially the same conformation in the dimeric structure with average RMSDs of 0.32 Å and 0.56 Å for the C _α_ atoms of the α and β subunit, respectively (Table S1). Chain B shows a slightly higher flexibility compared to that of chain A, which is reflected in a higher number of unmodeled residues (Table S2). A chloride ion was found in the catalytic site of the enzyme in the same position as it was in the truncated Taspase1 protein reported by Khan *et al*.(14) (Figure S2A), and an additional chloride ion was identified between the two protein monomers (Figure S2B). Also, a sodium ion was modeled just behind the active site (Figure S2C). The presence of Na^+^ ions in similar locations have been reported in the literature for type 2 asparaginases, not having any influence in the function of these enzymes.

### Structure determination of the active cp human Taspase1

While the crystallization of the cp-Taspase1_α41-183/β_ construct was rather straightforward, the fully active, circularly permuted Taspase1 (cp-Taspase1_α41-233/β_) form of the enzyme was very difficult to crystallize. Highly purified enzyme was subjected to large-scale, high throughput screening and optimization procedures to grow crystals of sufficient size and quality for X-ray structure analysis. After anisotropy correction (see Materials and Methods section for details), the crystal structure of the catalytically active cp-Taspase1_α41-233/β_ protein was determined to 3.04 Å resolution by molecular replacement using the structure of the inactive circularly permuted cp-Taspase1_α41-183/β_ protein as initial search model for phasing. The final structure was refined to R_work_ and R_free_ values of 25.5% and 31.7%, respectively.

Crystals of cp-Taspase1_α41-233/β_ protein contained the dimer of the α and β subunits (or hetero-tetramer) in the asymmetric unit with the typical αββα folding observed in this hydrolase family, in which the β strands are located in an antiparallel sandwich, and the α helices are located on the faces of the sandwich (Figure 3). All residues of the α and β subunits of both monomers have been correctly modeled except for a small loop (Ser352-Gln362), the last four residues of the long helix domain (Gly230, Thr231, Leu232, and Asp233), and the His-tag at the N-terminus. The structure of Taspase1 presented here has very good refinement statistics and excellent geometry (bond lengths and bond angles) as shown in Table 1.

**Figure 3.**
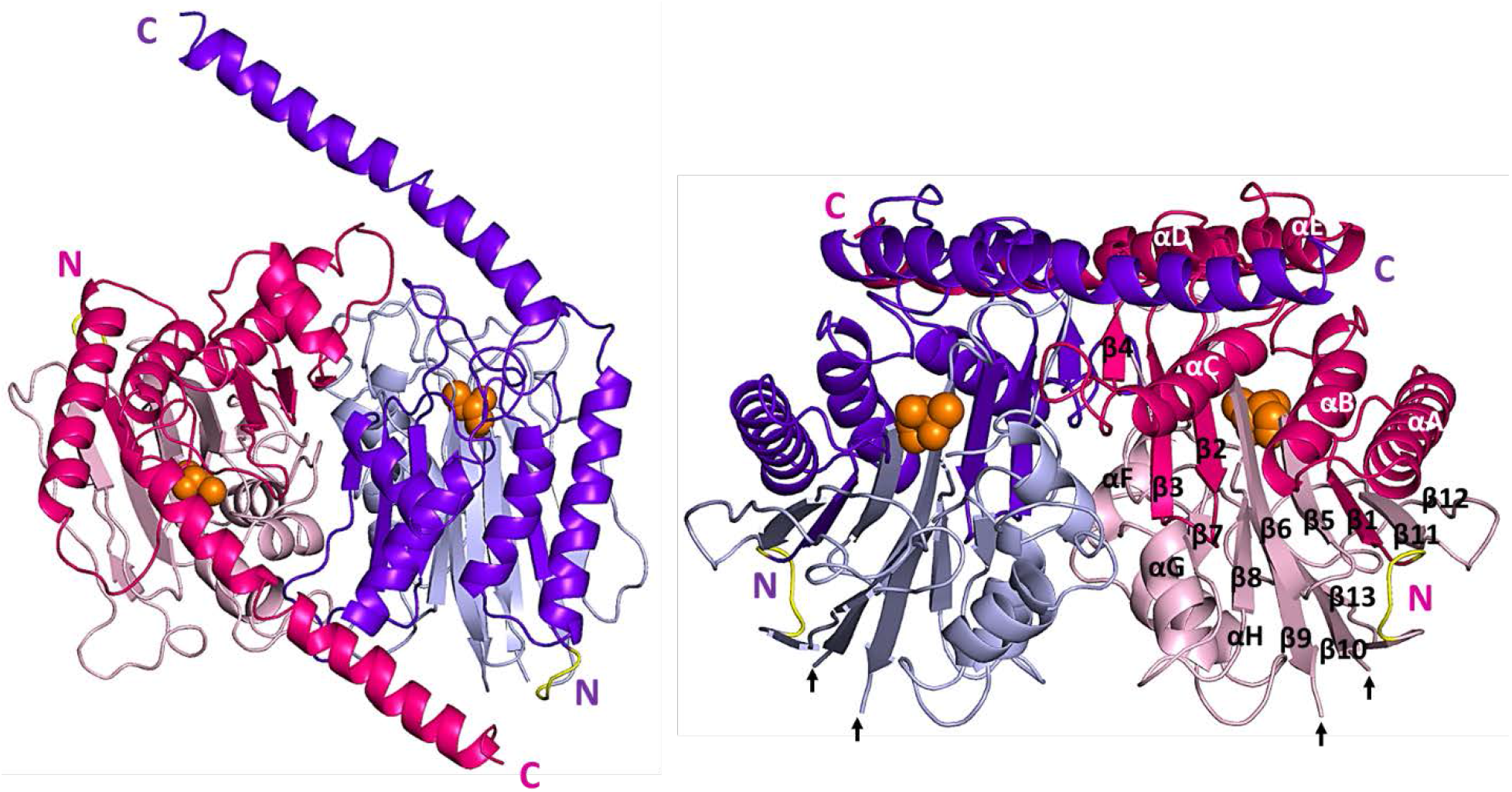
Crystal structure of cp-Taspase1_α41-233/β_. Side (left) and top (right) views of the hetero-tetramer. Cartoon representation of the α subunit (pink) and β subunit (light pink) of chain A and α subunit (purple) and β subunit (light blue) chain B of the dimer. All secondary structure elements as well as the N- and C-termini are indicated. The newly identified long helical domain (helix αE), points away from the dimer. The nucleophile Thr234 is show as orange spheres. The linker connecting the alpha and beta subunits is depicted in yellow. The missing fragment (Ser352-Gln362) is indicated by the arrows.

Despite the severe anisotropy featured by the crystals (Figure S3), the electron density maps of our structure show a high level of structural details. The 2mF_o_-DF_c_ electron density maps of the long helix domain (residues 183-228) can be used to assess the quality of our structure (Figure S4). Model-induced structural bias is absent in our Taspase1 structure, as positive electron density of the residues comprising the long helical domain was clearly visible already in the very initial stages of refinement (Figure S5). Two more regions from the core of the protein (the helical fragment from Ala58 to Ser96 and the beta strand fragment from Thr234 to Ser252) have been randomly selected to prove the substantial improvement of the electron density after anisotropy correction (Figure S6).

The most relevant finding of this new structure was the identification of the long helix, extending between Pro183 and Asp228, that is required for the activity of the enzyme. This segment was either absent in previous Taspase1 structures (14), or only partially resolved (Pro183-Lys202) in the proenzyme (14). Recently, van de Boom and co-workers proposed conformations for this long fragment before and after autoproteolytic cleavage based on molecular dynamic simulations, circular dichroism, and NMR (18) with the structure of the proenzyme as reference (PDB 2A8I (14)). While they predicted this domain to form a partially folded helix-turn-helix conformation (18), our structure instead reveals a novel conformation for this long fragment forming a long straight helix pointing outwards in opposite directions of the dimer (Figure 3). Similar to what was previously described by van de Boom and co-workers, the long helical fragment is interrupted by a turn-like element between residues Glu204 and Glu207 (Figure 3).

A structural comparison of cp-Taspase1_α41-233/β_ with previous Taspase1 structures shows that, while there is a high degree of similarity in the core of the protein, significant structural differences have been found in the loop areas and in the long domain that confers activity to the protein and was missing in all previous structures. Statistical results and RMSD values for the comparisons are in Table S3. A superposition of the active cp-Taspase1_α41-233/β_ structure to the published Taspase1 structures as well as to our inactive short cp-Taspase1_α41-183/β_ structure is shown in Figure S7. Interestingly, the long helix domain is in close proximity to the catalytic site, and slight but potentially important differences have been found in residues such as Tyr52 and Lys57 in the catalytic site (Figure 4). These differences become more pronounced when comparing cp-Taspase1_α41-233/β_ to the proenzyme. For instance, the side chain of Tyr52 is pointing toward the catalytic site in cp-Taspase1_α41-233/β_ and outward in the proenzyme. A comparison of the catalytic sites of chains A and B of the cp-Taspase1_α41-233/β_ with all Taspase1 structures is shown in Figure S8.

**Figure 4.**
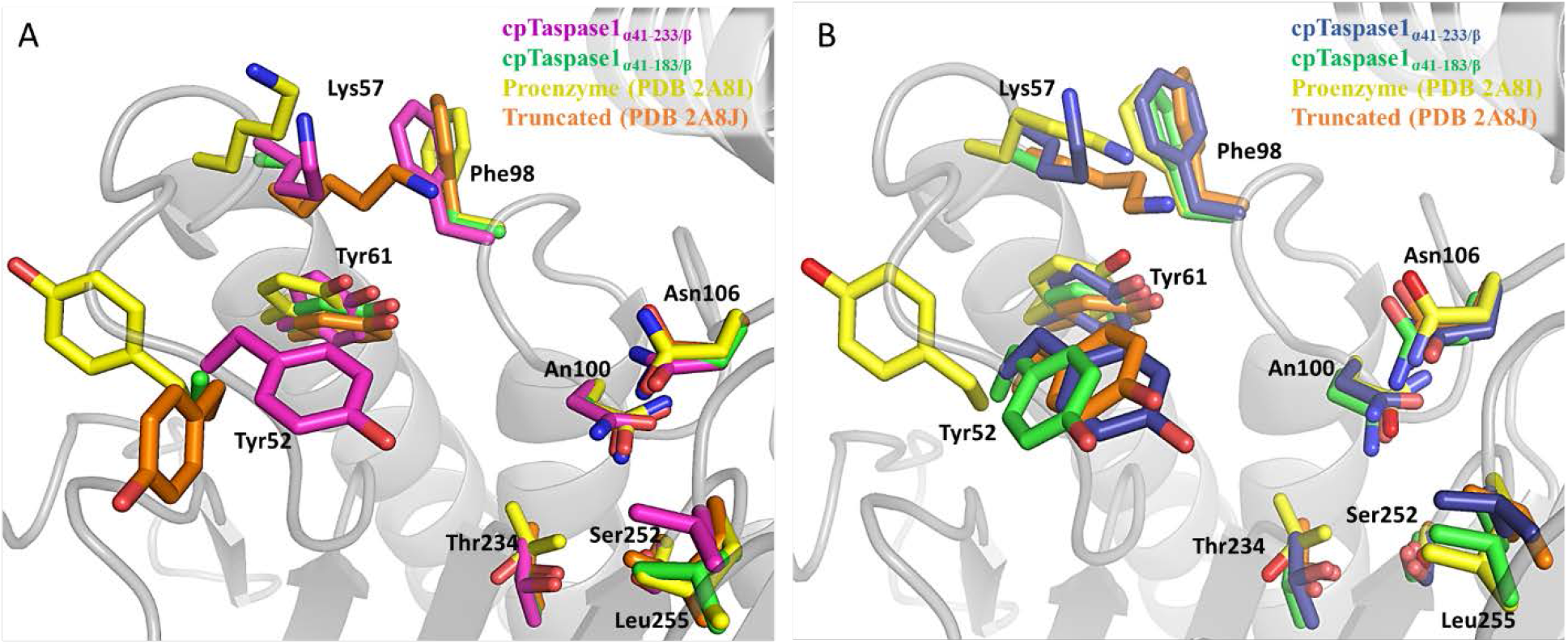
Comparison of the catalytic site of cpTaspase1_α41-233/β_ with other Taspase1 structures. Superimposition of chain A (A), and chain B (B) of cpTaspase1_α41-233/β_ to the proenzyme (PDB 2A8I (14)), the truncated Taspase1 (PDB 2A8J (14)), and the cpTaspase1_α41-183/β_. Residues important for the catalytic activity of Taspase1 are shown in stick representation.

As mentioned above, the largest structural differences are found at loop areas. One of those is the loop formed between Gly153 and Arg159, which has not been resolved in the previous mature, 2-chain truncated Taspase1 structure (14) and the cp-Taspase1_α41-183/β_ (this study), due to its high flexibility. In contrast, this loop is ordered in the structure of the active Taspase1 and has been fully modeled in our cp-Taspase1_α41-233/β_ structure. The tip of this loop of one of the monomers is slightly bent backwards and points towards the middle of the long helix domain of the neighboring monomer that stabilizes it (Figure S9). The loop fragment (Gly153-Arg159) was also fully modeled in the proenzyme (14), but has a slightly different conformation (Figure S9), as the amino acids of the helical domain from Leu203 onward were flexible and thereby could not be resolved in the structure of the inactive proenzyme.

Taken together, the change of the conformation of Tyr52 in the catalytic site and the bent conformation of the loop from Gly153 and Arg159 along with the long helix domain itself make the catalytic site of the protein adopt a narrower and deeper conformation than that of the truncated Taspase1 structures and the proenzyme (Figure S10). The calculated accessible surface area (ASA) of the catalytic site for the active cp-Taspase1_α41-233/β_ is larger than those calculated for the truncated proteins (cp-Taspase1_α41-183/β_ and Taspase1_α41-206/β_ (PDB 2A8J (14)) and the proenzyme (Table S4).

### Overall structure of the active cp human Taspase1

Crystals of the cp-Taspase1_α41-233/β_ protein belonged to the space group H32. An extended view of the asymmetric unit reveals that the αββα hetero-tetramer is part of a larger assembly, a double-ring like structure. An overview of the crystal packing of the cp-Taspase1_α41-233/β_ protein showing the rearrangement of the double-ring assembly is illustrated in Figure S11. This packing is completely different from what has been so far observed for all the truncated forms of the enzyme or the proenzyme, in which each single-ring forms a trimer of αββα hetero-tetramers, with the long helix domain being disposed sequentially in an up-and-down configuration. A schematic overview of the arrangement of the αββα hetero-tetramers in the single-ring assembly is shown in Figure 5A-C along the crystallographic ternary axis. In this arrangement, the three αββα units are identical, where the beta subunits are located closer to the 3-fold axis and form tight interactions within the trimer, while the alpha subunits are located at the periphery. As shown in Figure S12A showing the side view of the double-ring structure, the single-ring structure is arranged as layers that are mostly connected through the long alpha helix. Analysis of the dimer interfaces that form the double-ring assembly carried out by PDBePISA (https://www.ebi.ac.uk/pdbe/pisa/)server (19) revealed that the double-ring structure is stabilized through seven highly symmetric interfaces. Information about all the interfaces is listed in Table 2. All interfaces are shown in Figures 5D and S12B. A careful analysis of the interfaces is described in the Supplementary Information material.

**Table 2.** Interfaces in Cp-Taspase1_α41-233/β_ calculated by PDBePISA(19)

| Interface # | Name of Interface | Interface residues component 1 | Interface residues component 2 | BSA* (Å<sup>2</sup>) | Hydrogen bonds | Salt bridges | ΔG Kcal/mol | CSS<sup>#</sup> |
| --- | --- | --- | --- | --- | --- | --- | --- | --- |
| 1 | Dimer | 60 | 59 | 2039.1 | 21 | 7 | -23.8 | 1.000 |
| 2 | Inter-single ring | 42 | 43 | 1355.1 | 13 | 2 | -9.5 | 1.000 |
| 3 | Helix | 19 | 19 | 648.2 | 2 | 0 | -6.6 | 0.069 |
| 4 | Intra-double ring | 16 | 16 | 535.7 | 0 | 0 | -9.7 | 0.000 |
| 5 | Linker | 8 | 8 | 168.0 | 0 | 2 | -1.7 | 0.026 |
| 6 | Histidines | 1 | 1 | 36.1 | 0 | 2 | 0.1 | 0.006 |
| 7 |  | 1 | 1 | 33.6 | 0 | 2 | 0.1 | 0.006 |

\*Buried surface area (BSA).
<sup>#</sup>Complex formation significance score (CSS).

**Figure 5.**
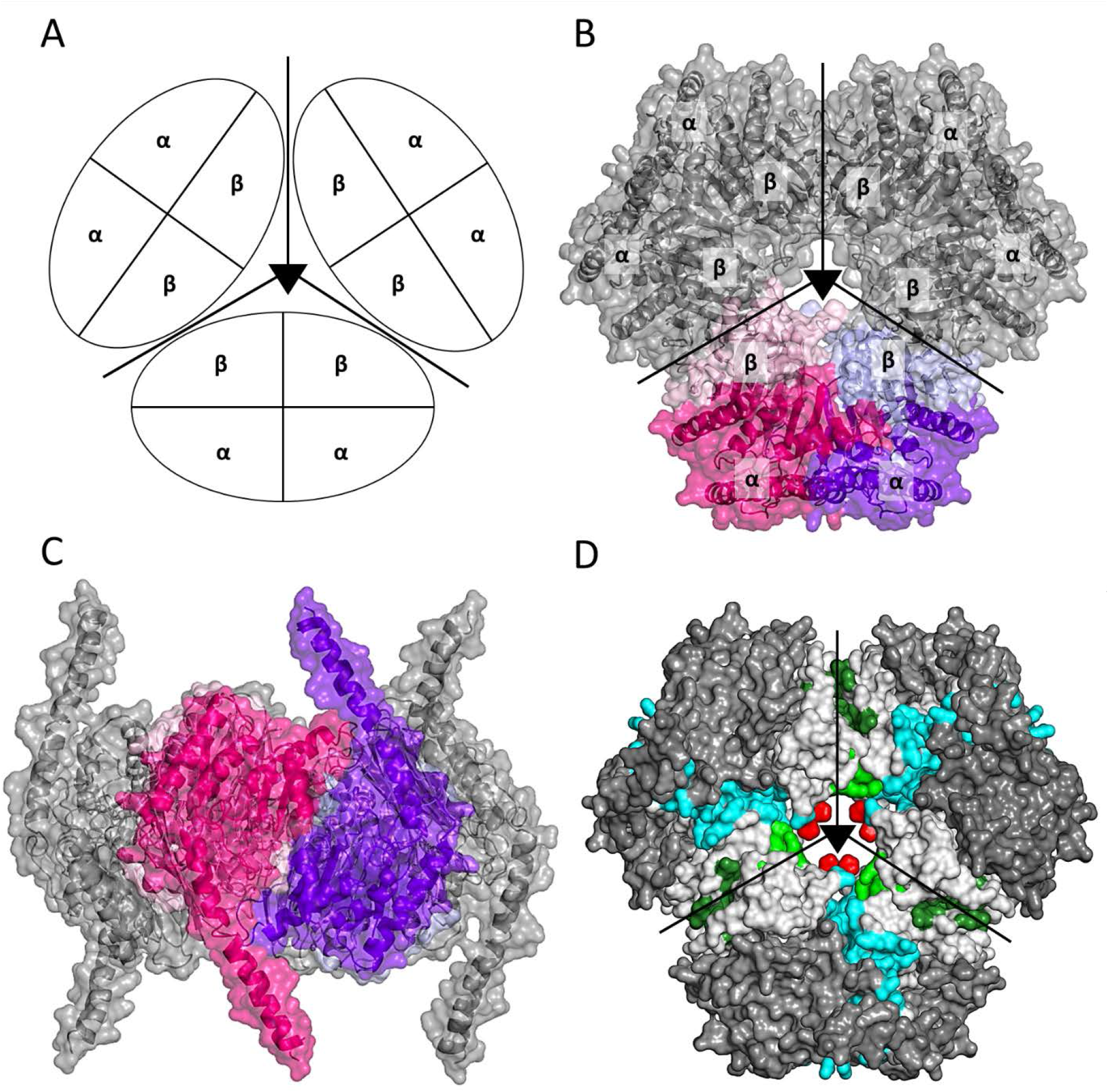
Subunit interfaces that maintain the single-ring assembly of cpTaspase1_α41-233/β_ in the crystals. A) Schematic view of the single-ring structure. B) Top view of the surface representation of the cpTaspase1_α41-233/β_ of the single-ring structure. C) Side view of the surface representation of the cpTaspase1_α41-233/β_ of the double-ring structure D) Interfaces at the single-ring structure. The interfaces are colored as follows: cyan for the dimer interface, green for the intra-ring interface, and red for the histidine interface. The cpTaspase1_α41-233/β_ dimers are represented with the same color as in Figure 3 and the dimers generated by symmetry are shown in a grey and black.

### DLS confirms a concentration dependent oligomeric state

To study whether the single and/or double-ring structures observed in the crystals exist in solution or are a crystallization artifact, we performed DLS experiments with Taspase1 at various concentrations (2.5, 5, and 10 mg/ml) in the purification buffer in the absence of crystallization agents. As shown in Figure S13 and S14, a uniform and narrow size distribution was observed for all samples, indicating that only one species was present in solution at each concentration. However, the hydrodynamic radius of the observed species increased from ∼5 nm to approximately ∼5.5-5.6 nm as the protein concentration increases. Those hydrodynamic radii were estimated to correspond to protein species between 150 kDa and ∼225 kDa, indicating that the oligomeric state of Taspase1 is strongly concentration-dependent varying from tetrameric at 2.5 mg/ml to hexameric at 10 mg/ml. This behavior suggests that different oligomeric forms of Taspase1 may exist in solution, and possibly, also in cells.

### EM suggests the dimer as predominant assembly in solution

To further examine the oligomeric state of the protein using an alternative method, we performed negative staining electron microscopy experiments. The electron micrographs show a highly uniform distribution of particles, with an estimated size of ∼7-8 nm (Figure 6A). The sample was further analyzed by using 2D class averaging of negative staining EM micrographs. We identified a total of 250 particle classes, three of which are shown in Figure 6B. The sizes of all those particles was estimated to be ∼7.5 nm. Thus, even at extremely low concentrations, Taspase1 forms large particles that resemble in both shape and size those of the αββα hetero-tetramer (Figure 6C). Interestingly, the orientation of some of the particles allowed us to identify the long domain. However, no evidence of double-ring structures was found, which indicates that the higher assembly is an artifact of crystallization.

**Figure 6.**
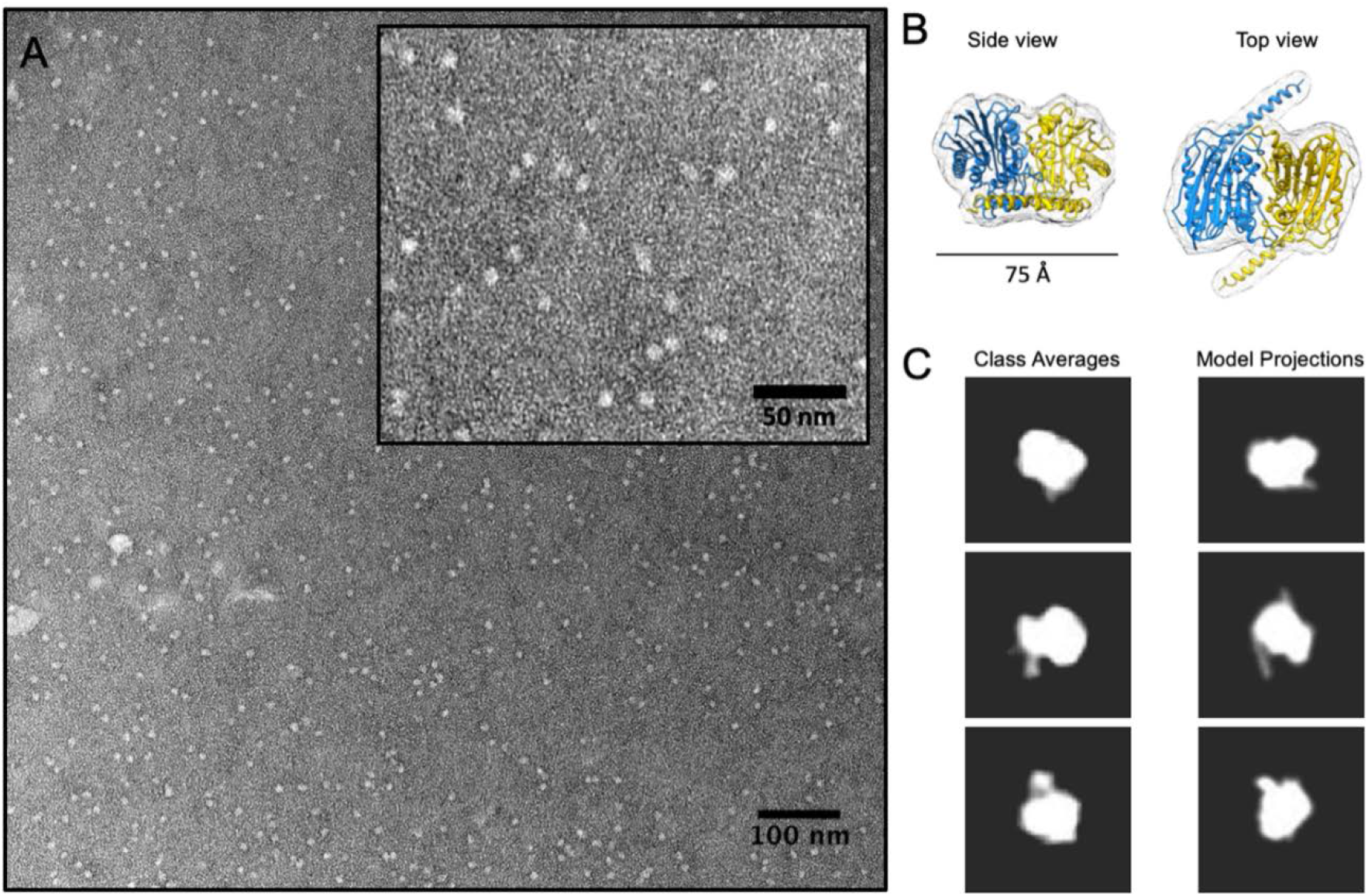
Negative stain electron microscopy (EM) of cpTaspase1_α41-233/β_. A) EM micrograph showing a distribution of cpTaspase1_α41-233/β_ particles. Inset panel shows a closer view of the particles. Particles were estimated to be ∼6-7 nm in size. B) Model of the cpTaspase1_α41-233/β_ dimer that was used to generate 15Å density map (shown as mesh around the model). C) Projection images were generated using the calculated map in B) and these were compared with 2D class averages calculated from the negative stain images collected from cpTaspase1_α41-233/β_. Representative class averages and projection images of the model are shown, and it can be seen that he dimeric model matches well with the experimental class averages.

### Disorder prediction and coiled coil structure analyses

The propensity of Taspase1 to harbor intrinsically disordered regions (IDRs) was predicted using four different web servers: PONDR-FIT (20), MetaDisorder (21), MobiDB (22), and SPOT-Disorder2 (23). All four tools predicted the long domain (Pro183-Asp233) of Taspase1 to be highly disordered (Figure 7A). According to PONDR-FIT and MetaDisorder, this could be a structural disorder, as it encompasses at least 25 or more consecutive residues of the helix (Figure 7B and Table S6). Further analysis with the Protparam-Expasy tool (24), showed that the long domain is enriched in charged and flexible residues (disorder-promoting residues) such as R, S, E, D and K, as well as lack/low abundance of hydrophobic residues (order-promoting residues) such as G, C, P, V, W and Y, which explains the high tendency of this fragment to be disordered (Figure 7C). The long domain was also inspected by several coiled coil structure predictors including COILS (25), DEEPCOIL (26), and MARCOIL (27). All three predicted the long fragment to have a high tendency to form a coiled coil structure (Figure S15).

**Figure 7:**
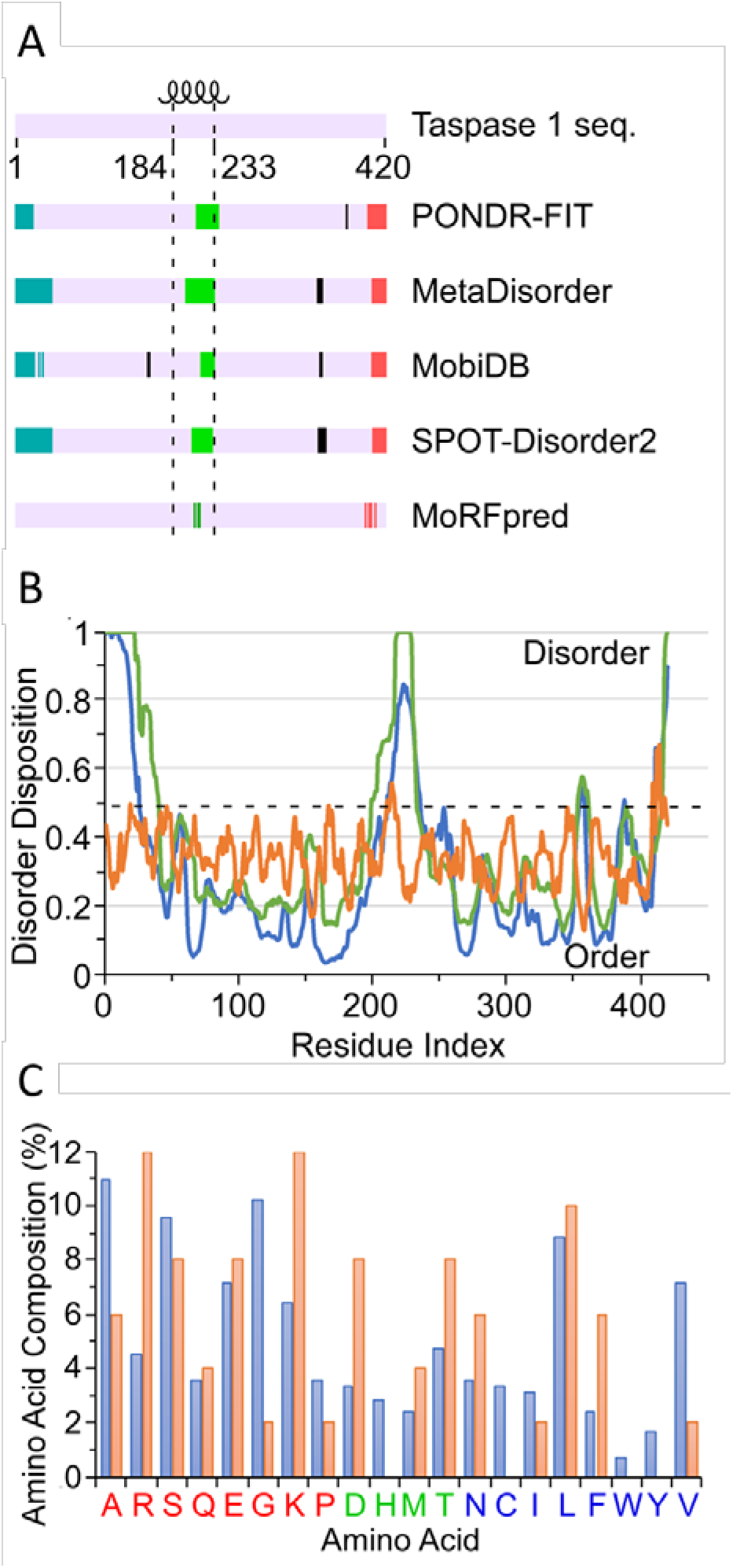
Prediction of intrinsically disordered regions (IDRs) of Taspase1. A) Disorder predictions in three discrete regions of Taspase1 (N-terminal (cyan), middle (green) and C-terminal (red)) by PONDR-FIT (20), MetaDisorder (21), MobiDB (22), MoRFpred (40), and SPOT-Disorder2 (23). Short disordered regions are represented in solid black. The location of the long helix fragment of Taspase1 is delimited by the black dotted line. B) Disorder disposition of each residue in Taspase1 sequence predicted by PONDR-FIT (blue), MetaDisorder (green) and MoRFpred (dark orange). Residues with disorder probability higher than 0.5 (black dotted line) are predicted to be disordered. C) Comparison of the composition of each amino acid in the long helix (coral orange) with full-length Taspase1 (blue). Disorder-promoting residues are red, order-promoting residues are blue and disorder-order neutral residues are green.

## Discussion

In this study, we have determined the 3D structure of the fully active form of human Taspase1 to 3.04 Å resolution, capturing the protein with a previously unobserved, important structural feature, a long C-terminal domain presented as a nearly continuous helix projecting outward from the dimer, which is essential for enzymatic activity. Previous molecular dynamic simulations and NMR methods suggested that this piece of the protein (Pro183-Asp233) forms a helix-turn-helix conformation and adopted two different states, a closed state in the proenzyme and an open state that is acquired upon autoproteolytic cleavage (18). Yet, little is known about this newly discovered long helix domain of Taspase1 or the structurally related type 2 asparaginases. Despite its proximity to the catalytic site, this portion of the protein had not been thought to be important for catalytic activity, perhaps because its mobility had prevented it from being observed in the crystal structures previously reported (10-14, 28-36). While it has been demonstrated that mutations within the catalytic site of type 2 asparaginases, including Taspase1 (14), hASRGL1 (37), glycosylasparaginase (38), and hASNase3 (39), and others, abolish or significantly diminish the catalytic activity of the proteins, an essential role for a segment like the Pro183-Asp233 helix of Taspase1 had not previously been considered. In addition to cp-Taspase1_α41-233/β_, we have generated two shorter protein variants, the cp-Taspase1_α41-206/β_ containing half the helix (Pro183-Asp206), and the cp-Taspase1_α41-183/β_, which lacks the entire fragment. While cp-Taspase1_α41-233/β_ was fully active the catalytic activity was diminished in cp-Taspase1_α41-206/β_ and completely abolished in cp-Taspase1_α41-183/β_. These results suggest that a similar behavior could be expected for the long fragment of structurally-related type 2 asparaginases in which the full length of the helical domain may be needed for the enzymes to develop their function.

Mutational studies of residues in the catalytic site (14) led to the currently accepted model for the proteolytic mechanism of Taspase1 in which a small set of amino acid residues (e.g., the nucleophile Thr234) are considered important to the catalytic reaction. However, we have demonstrated that the loss of the long domain (partially or fully) dramatically affects the activity of Taspase1 and thus raises questions as to the critical role of this region in the function of Taspase1. We have also observed that the presence of the long domain leads to conformational changes, specially Try52 and the loop Gly153-Arg159, which could be somewhat implicated in the catalytic mechanism of Taspase1. Those conformational modifications create a narrower and deeper catalytic site, and possibly could help stabilize and facilitate substrate binding. In the case of the Tyr52, even though is far from the active site, it could contribute to stabilize and redirect the substrates into the catalytic site by interacting with them. Thus, the most likely role for the long domain in the proteolytic cleavage of substrates by Taspase1 would be to facilitate substrate binding. However, we currently have no experimental evidence, such as structures with bound substrate or product, that might support this proposal. Having crystal structures of cp-Taspase1_α41-233/β_ protein in complex with substrates is absolutely required to fully understand how Taspase1 recognizes and processes its targets. We therefore conclude that the current proteolytic mechanism of Taspase1 and, overall, the type 2 asparaginases family must be revised, and the long domain, which has been largely ignored so far, must now also be considered as a key part of the mechanism.

The observed straight helical conformation that points outwards and interacts with another long helix of a neighboring molecule resembles that of a typical coiled coil-type structure. This novel feature has been furthered supported by sequence analyses where this fragment was predicted to have a high tendency to form coiled-coil type structures. In the recent study by van de Boom and co-workers (18), the long fragment was predicted to form two helices separated by a short turn (a helix-turn-helix conformation), in which the two helices interact with each other to adopt a folded conformation typical of coiled-coil structures (18). Additionally, we have shown that the long domain of Taspase1 has also been predicted to be intrinsically disordered using various disorder predictors algorithms, which is consistent with the fact that the long domain has been unresolved in all Taspase1 structures reported so far. We also have used MoRFpred (40) to predict that molecular recognition features (MoRFs) within the long-disordered region. MoRFs are short functional IDRs located in long disordered regions and undergo disorder-to-order transition upon interacting with their partners (41). MoRFpred predicted several MoRF residues within both middle- and C-terminal-disordered regions (Figure 8A, B; and Table S6) of Taspase1. Disordered regions in proteins have often been associated with function (42-45). A common characteristic of disordered regions is their ability to become ordered, typically leading to the formation of α helices, upon binding of substrates from natural targets (44).

Taken together, the long domain of Taspase1 might adopt a coiled coil-type structure promoted by its natural substrates. Taspase1 has been shown to contain a highly evolutionary conserved bipartite NLS at the long domain between residues 197 and 220 (^197^<u>KR</u>N<u>KRK</u>LELAE<u>R</u>VDTDFMQL<u>KKRR</u>^220^), which could be utilized to translocate the protein into the nucleus by direct binding to importin-α. Therefore, a plausible hypothesis is that Taspase1, initially expressed as an inactive precursor, could use its long domain to translocate from the cytoplasm to the nucleus, where ultimately undergoes intramolecular cleavage to become the active protease, through the interaction of its nuclear localization sequence (NLS) with importins. Importins α possess a highly structured helical domain comprised of ten tandem armadillo (ARM) repeats (46-48), whose interior serves as the NLS-binding groove by wrapping it in a fashion that resembles a twisted slug (49-51). The structure of the cp-Taspase1_α41-233/β_ protein reported in this study provides the first structural visualization of the NLS within the context of the Taspase1 protein. We can, therefore, propose that the long domain of Taspase1 binds to importin α in a similar fashion to that found for the NLS of the Fused in Sarcoma (FUS) protein (49-51). The NLS of FUS protein, which is initially intrinsically disordered, becomes ordered by adopting a helical conformation upon binding to importin α. Taken together, we hypothesize that, as observed for FUS protein, the binding of Taspase1 to importin α may trigger large conformational changes in the initially disordered long domain inducing it to become the long straight helix observed in our crystals.

Controversy concerning the functional multimeric state of Taspase1 exists. All structural studies reported so far support the formation of the hetero-tetramer, in which the αββα subunits remain together upon autoproteolytic cleavage (14), which was postulated by others based on the high homology of Taspase1 with the Type 2 asparaginase and the Ntn hydrolase families (3, 4, 52). However, studies carried out in living cancer cells suggested Taspase1 functions as an αβ monomer (53, 54). The approach pursued by Bier *et al*. examined whether interference of Taspase1 dimer formation via expression of dominant-negative mutants could impede Taspase1 from processing its natural targets. However, this strategy, that has been successfully applied to other systems (55-59), was shown to be not applicable to Taspase1 as the activity was not inhibited when WT Taspase1 was in contact with inactive mutants (53). Bier, *et al*. argued that this outcome is due to the failure of Taspase1 to form the αββα hetero-tetramer *in vivo*. More recently, Sabiani and co-workers proposed a model for the proteolytic mechanism in which the hetero-tetramer formation process was suggested to be the key event that triggers the activation (or maturation) of Taspase1 (60). The results of Bier and co-workers argue against the hetero-tetramer formation. However, in their studies, Taspase1 was expressed as a fusion protein (i.e., with GFP or mCherry) to monitor fluorescence and both fusion tags are over half the size (∼27 kDa) of Taspase1 (50 kDa). Thus, it is possible that hetero-tetramer formation might be impeded. In addition to this, Bier and co-workers argue that the Khan *et al*.’s crystallographic results show that the αββα structure is a crystallization artifact (53). However, Khan *et al*. co-expressed and purified the individual α and β subunits, leading to the αββα hetero-tetramer later seen in their crystallization studies. Also, as shown in our results, the hetero-tetramer interface buries a large surface area that is stabilized by numerous contacts (see Table 2 and S5). Large and stable interfaces such as these are unlikely to result only from crystal contacts but are more likely to be interfaces forming functional oligomeric states. Our DLS and EM studies also demonstrate that the smallest assembly of Taspase1 in solution is the αββα hetero-tetramer, confirming what has been proposed by published Taspase1 structures (14). DLS also demonstrated that the oligomeric state of Taspase1 is concentration dependent, forming higher oligomeric states -plausibly tetramers and hexamers- in solution at higher protein concentrations. We therefore hypothesize that the biological multimeric state of Taspase1 is the αββα hetero-tetramer that eventually self-oligomerizes to form a higher oligomeric assembly in the form of a trimer of αββα hetero-tetramers, leading to the single-ring structure. Further *in vitro* and *in vivo* studies, as well as structural analysis, will be needed to understand how and why Taspase1 oligomerizes and what are the biological functions or consequences of this self-oligomerization.

Taspase1 is classified as a “non-oncogene addiction” protease overexpressed in primary human cancers and its deficiency disrupts proliferation of human cancer cells *in vitro* and in mouse tumor xenograft models of glioblastoma. Taspase1 is, therefore, considered a novel anticancer drug target; however, to date the only reported inhibitors are an arsenic acid-based small molecule (NSC48300) and succinimidyl-containing peptides with modest potency (61),(62). In order to advance the design of Taspase1 inhibitors for cancer therapy, more structural information in combination with biochemical and cellular assays to specifically assess inhibition of Taspase1 will be needed. In this regard, the structure of Taspase1 reported in this study, along with the activity assay, offers new insights into the requirements for Taspase1 activity. Our results demonstrate that, even though residues essential for catalytic activity lie within the presumed catalytic site of Taspase1, another region of the protein strongly influences the catalytic activity of Taspase1, which was previously unappreciated. Further investigation of the role of the long domain in the catalytic activity of Taspase1 may offer opportunities for drug targeting and suggest potential alternative routes to inhibition.

## Materials and Methods

### Cloning, expression, and purification of the circularly permuted human Taspase1 proteins

When examining the structure of the proenzyme Taspase1 (PDB 2A8I) (14), the proximity of the C-terminus of the C-terminal domain (CTD) to the first AA observed in the N-terminal domain (NTD), G41, was noted. This juxtaposition suggested that insertion of a short linker (GSGS) between these ends might enable the creation of circularly permuted (cp), single chain constructs, which might simplify expression and purification of different forms of Taspase1. Three circularly permuted constructs (cpTaspase1_α41-183/β_, cpTaspase1_α41-206/β_ and cp-Taspase1_α41-233/β_) were designed. Expression and purification of each of these proteins has enabled us to characterize their enzymatic activity and determine novel crystal structures.

The expression vector pEMB7013 harboring the sequence of the cp-Taspase1_α41-233/β_ (a.a. 234-416-GSGS-41-233), the partially truncated cpTaspase1_α41-206/β_ (a.a. 234-416-GSGS-41-206) or the fully truncated cpTaspase1_α41-183/β_ (a.a. 234-416-GSGS-41-183) human Taspase1 constructs with a hexa-histidine tag at the C-terminus was transformed in the *E. coli* BL21 (DE3) strain. Transformed cells were grown at 37°C in 2xYT medium supplemented with 50 μg/ml kanamycin, and expression of Taspase1 was induced with 0.5 mM IPTG when the OD_600_ reached values between 0.5 and 0.7. Following growth overnight at 20°C, cells were harvested by centrifugation at 7,500 x g for 20 min at 4°C and stored at −80°C for purification later. Purification of the three proteins was carried out using the same procedure. Cell pellets were re-suspended in 50 mM Tris-HCl pH 8.0, 500 mM NaCl, 5% glycerol, 0.25 mM TCEP, 0.5% CHAPS, 1 mg/ml lysozyme and 30 U/ml Benzonase, supplemented with EDTA-free protease inhibitor cocktail (Sigma-Aldrich). Cells were lysed by sonication using 5 cycles in pulsed mode with 1 min rest on ice between cycles. Each cycle consisted of thirty 2 second (s)-pulses at 70% amplitude and cooling for 2s between pulses. The lysate was subsequently centrifuged at 45,000 x g for 1 h at 4°C to remove unbroken cells and cell debris. The supernatant was then filtered through 0.22 μm filters and loaded onto a Ni-NTA HisTrap™ HP column previously equilibrated with equilibration buffer (EB) 50 mM Tris-HCl pH 8.0, 500 mM NaCl, 5% glycerol, 0.25 mM TCEP, 10 mM imidazole. After collection of the flow through, the column was washed with 20 column volumes of EB followed by another wash with 15 column volumes of EB containing 25 mM imidazole. Taspase1 proteins were finally eluted with EB using a linear gradient of imidazole between 25 and 250 mM. Fractions containing Taspase1 protein were combined and concentrated to 12-15 mg/ml using 30 kDa cut-off concentrators. The protein was further purified by size exclusion chromatography (SEC) using a HiPrep™ 26/60 Sephacryl™ S-200 HR (GE healthcare) column previously equilibrated in 20 mM HEPES pH 8.0, 500 mM NaCl, 5% glycerol. Fractions containing Taspase1 protein were pooled, concentrated as described above, flash-frozen using liquid N_2_, and stored at −80°C for later use. All purification steps were carried out at 4°C using the ÄKTA Pure fast protein liquid chromatography (FPLC) system (GE healthcare). Protein purity was assessed by SDS-PAGE. Purification results of the cp-Taspase1_α41-233/β_ are illustrated in Figure S16A, B.

### Activity assays of the circularly permuted human Taspase1 proteins

To assess the catalytic activity of the designed cp Taspase1 constructs, a quenched fluorescence resonance energy transfer (FRET) based enzyme assay was performed (63) that measures proteolysis of a labeled Taspase1 substrate by the purified cp Taspase1 constructs. The substrate consisted of a 10-residues peptide derived from the second cleavage site of the endogenous Taspase1 substrate MLL, flanked by a MCA (7-Methoxycoumarin-4-yl)acetyl) fluorophore and 2,4-dinitrophenyl-lysine (Lys(DNP)) as the quencher: MCA-Lys-Ile-Ser-Gln-Leu-Asp↓Gly-Val-Asp-Asp-Lys(DNP)-NH_2_. The peptide was purchased from CPC Scientific Inc (San Jose,CA). In the absence of Taspase1, the fluorophore does not emit due to close proximity of the quencher. The proteolysis of the peptide substrate by Taspase1 releases the quencher from the fluorophore and generates a fluorescence signal. A schematic of the proteolytic reaction is illustrated in Figure S17. Fluorescence intensities were measured with excitation 340/emission 455 filters using a Perkin Elmer Envision multi-mode plate reader. The reaction buffer contained 100 mM ammonium acetate pH 8.3, 625 µM β-mercapto-ethanol (BME), 10 μM substrate, and 100 nM enzyme.

Taspase1 enzymes were tested for activity by a kinetic read method in triplicate alongside a “substrate only” negative control in a 384 well plate. Buffer (5 µl) and enzyme (10 µl) were added to each well at final concentrations of 100 mM NH4Ac pH 8.3, 625 µM BME plus 100 nM Taspase1 enzymes, except for the “substrate only” wells that received 10 µl of additional assay buffer instead of enzyme. The reactions were started by adding 10 µl of 10 µM MCA-DNP substrate to all the wells. Plates were read on the EnVision multi-mode plate reader, which was immediately initialized to generate reaction progress curves. Readings were taken every minute for 90 min at room temperature.

### Crystallization of the circularly permuted human Taspase1 proteins

Automated, high-throughput crystallization experiments were performed using various commercial kits. Initial crystallization conditions of truncated cp Taspase1 (cpTaspase1_α41-183/β_) at ∼9-10 mg/ml were identified from PACT screen (B11) condition (0.2 M calcium chloride, 0.1 M MES pH 6, 20% (w/v) PEG 6000). For cpTaspase1_α41-233/β_, initial hits were obtained and further optimized with the best crystals of cpTaspase1_α41-233/β_ being obtained at 20°C using hanging drop vapor diffusion method. Protein at 15 mg/ml was mixed in a 2:1 ratio with precipitant solution composed of 0.1 M sodium citrate pH 4.0, 1.0 M ammonium sulfate, 10% 2-methyl-2,4-pentanediol (MPD), and 0.4 M NaCl. Crystals in these conditions grew in 3 days and reached sizes between 200 and 500 µm in their longest dimension (Figure S16C).

### Data collection, structure determination, and refinement of the cpTaspase1_α41-183/β_ protein

Crystals of cpTaspase1_α41-183/β_ grown in the above conditions were cryo-protected using 20% ethylene glycol in the crystallization buffer. Data sets were collected at beamline 21-ID-F at the Advanced Photon Source (APS), with a RAYONIX MX-300 detector at a wavelength of 0.97872 Å under a stream of nitrogen (100K). Data were indexed and integrated with XDS/XSCALE (64). The structure was solved by molecular replacement with MOLREP (65) using the structure of the mature, 2-chain truncated Taspase1 (PDB 2A8J(14)) as search model. The final structure was solved at a resolution of 2.15 Å after numerous iterative rounds of manual rebuilding in COOT (66) and refinement in PHENIX (67). The structure was examined, validated, and improved using MolProbity. The crystallographic data collection and structure refinement statistics are summarized in Table 1.

### Data collection, structure determination, and refinement of the cpTaspase1_α41-233/β_ protein

For data collection, crystals grown in the conditions above indicated were transferred into a cryoprotectant solution of 30% glycerol in the crystallization buffer for less than 5 seconds before being flash cryo-cooled in liquid nitrogen. X-ray data sets were collected for the cpTaspase1_α41-233/β_ crystals at 100 K at the APS GM/CA 23-ID-D beamline using a Pilatus 16M detector. The diffraction images were indexed and integrated with XDS (64) in H32 space group with unit cell dimensions of a=196Å, b=196Å, c=196.91Å. The dataset was scaled, and the structure factors generated using AIMLESS (68) and TRUNCATE (69), respectively. Using XTRIAGE program (70) in PHENIX suite (67), the crystals were confirmed not to be merohedrally twinned.

Phasing was successfully performed by molecular replacement with PHASER (71) using the structure of the circularly permuted truncated Taspase1 (cpTaspase1 _α41-183/β_) as the search model upon removal of all solvent molecules. Phaser found a unique solution (TFZ 9.5 and LLG 434) with two monomers in the asymmetric unit consisting of the hetero-tetramer a s previously reported for the crystallographic structures of the proenzyme and the truncated Taspase1(14). The structure was built and refined using REFMAC5 (72) and COOT (66). Extra positive electron density was clearly visible for the long fragment (Pro183-Asp233), that was missing in all previous structures, which was built and refined in later steps. As our crystals showed anisotropic diffraction, the initial refinement generated moderate refinement statistics, with R_work_ and R_free_ values of 32% and 36%, respectively, as well as poor electron density maps.

As shown in Figure S3A, the diffraction was highly anisotropic with the useful data extending only to a modest-low resolution (Table 1). Thus, structure determination of the cpTaspase1_α41-233/β_ was challenging. The TRUNCATE program (69) revealed two strong-diffracting directions (a* and b*) and one very weak-diffracting direction (c*) (Figure S3B). Anisotropy analysis was addressed with the STARANISO server (http://staraniso.globalphasing.org/cgi-bin/staraniso.cgi)using the DEBYE and STARANISO programs to perform an anisotropic cut-off of merged intensity data, to perform Bayesian estimation of structure amplitudes, and to apply an anisotropic correction to the data. The diffraction images were re-processed with XSCALE/XDS (64) with no resolution cut-off applied; an *ahkl* output file containing unmerged and unscaled intensities was generated and was provided as input for the STARANISO server using the unmerged STARANISO protocol, which prevents the noisy measurements in the weakly diffracting directions from adversely affecting the scaling and error-model estimation at the final scaling/merging step, using the default cut-off level of 1.2 for the local I/σ<I>. STARANISO indicated that the data were strongly anisotropic (the anisotropic ratio and S/N ratio were 1.474 and 121.60, respectively), which yielded an ellipsoidal resolution boundary with resolution limits of 2.9 Å, 2.9 Å, and 5.3 Å along three principal component a*, b*, and c* axes, respectively. After the anisotropic correction, the final dataset showed a best-resolution limit of 3.04 Å along the a*, b* axes, and a lowest-resolution limit of 5.8 Å along the c* axis. The axes-dependent truncated and corrected structure factors generated by STARANISO were then used to repeat the molecular replacement phasing using the initially refined model that resulted in significantly better solution (TFZ 37.1 and LLG 1070). This new model was refined with REFMAC5 with non-crystallographic symmetry (NCS) using local restraints. Density modification with PARROT program (73) using NCS averaging and solvent flattening was also used. The final model was refined to 3.04 Å resolution with R_work_ and R_free_ values of 25.5% and 31.7%, respectively. Moreover, the final model yielded electron density maps that were much more interpretable. The crystallographic data collection and structure refinement statistics are summarized in Table 1.

### Structure alignments

The alignments between crystallographic structures were carried out by LSQKAB using the method described by Kabsch (74). The root-mean square deviations (RMSDs) excluding the long domain residues (Pro183-Asp233) were calculated for the Cα and all atoms.

### Bioinformatics analyses

The sequence of the full length human Taspase1 (UniProtKB accession number Q9H6P5) was further analyzed. Intrinsic Disorder analysis was performed using five web-based disorder prediction tools: PONDR-FIT (20), MetaDisorder (21), MoRFpred (40), SPOT-Disorder2 (23), and MobiDB (22). Expasy-Protparam was used to calculate the frequency of occurrence of different amino acids (https://web.expasy.org/protparam/). Coiled-coil structural predictions were performed using three available predictors from EXPASY server: COILS (25), DEEPCOIL (26), and MARCOIL (27).

### Negative staining electron microscopy

Freshly purified cpTaspase1_α41-233/β_ protein was negative stained and examined by transmission electron microscopy (TEM) to assess cpTaspase1_α41-233/β_ protein homogeneity. Taspase1 at 0.52 μg/ml in the purification buffer (20 mM HEPES pH 8.0, 500 mM NaCl, 5% glycerol) was applied as 5 μl drops for 2 min at room temperature to carbon-coated 400 mesh copper grids, which were previously glow-discharged for 60 seconds. The excess liquid was removed by blotting then replaced with a drop (5 µl) of stain 0.75% (w/v) pH neutralized uranyl formate solution. Excess staining solution was removed promptly with filter paper and the staining step was repeated once more. Grids were air-dried at room temperature prior to data acquisition. Taspase1 particles were initially examined with a Philips CM12 transmission electron microscope operated at 120 kV. A second round of negative staining experiments was performed to obtain 2D-class averages. For these experiments Taspase1 was used at 1.6 μg/ml and examined with FEI Titan/ETEM microscope on a Gatan US1000XP CCD camera. A total of 69 micrographs were acquired with a 2.5 x 2.5 Å pixel and processed within the cisTEM software package (75). A total of 40,365 particles were used for statistical analysis and a total of 250 classes were generated and averaged in the cisTEM software suite.

### Dynamic light scattering (DLS)

DLS measurements were performed on two different instruments for comparison, a SpectroSize 300 spectrometer instrument (Molecular Dimensions), and a DynaPro Nanostar instrument (Wyatt Technology). Samples of cpTaspase1_α41-233/β_ were prepared at three different concentrations (2.5 mg/ml, 5 mg/ml, and 10 mg/ml) in in the purification buffer (20 mM HEPES pH 8.0, 500 mM NaCl, 5% glycerol). Immediately before measurements protein solutions were centrifuged for 30 min at 14000 rpm, 4°C, in order to remove any aggregates and dust. Instruments software were used in data collection and processing. All measurements were carried out by averaging 10 accumulations of 20 sec each at 20 °C.

## Data Availability

The structure models and structure factors for the cp-Taspase1_α41-233/β_ and cp-Taspase1_α41-183/β_ have been deposited at the wwPDB under accession codes PDB 6VIN, and PDB UGK, respectively.

## Supplementary information

The supplementary information includes a detailed description of all the interfaces found in the crystallographic structure of cp-Taspase1_α41-233/β_, Figures S1 to S23, and Tables S1 to S6.

## Acknowledgements

This project has been funded in whole with Federal funds from the National Cancer Institute, National Institutes of Health, under Chemical Biology Consortium Contract No. HHSN261200800001E. The content of this publication does not necessarily reflect the views or policies of the Department of Health and Human Services, nor does mention of trade names, commercial products, or organizations imply endorsement by the U.S. Government. We would like to recognize the protein expression, purification, and crystallization teams at Beryllium for their assistance. We thank Dr. Barbara Mroczkowski, Dr. Joel Schneider, and Dr. Michele Zacks, for their careful scientific editing and proofreading.

## Author contributions

J.M. Martin-Garcia, S. L. Delker, T. E. Edwards, P. Fromme, A. Flint, and J. Hsieh designed the experiments and provided with the most relevant scientific discussions. S. L. Delker and T. E. Edwards designed and cloned all proteins presented in this study. S. L. Delker and T. E. Edwards crystallized and determined the structure of the cp-Taspase1_α41-183/β_ protein variant. L. Sambucetti, M. Stofega, and J. Snider performed all catalytic activity assays of all proteins presented in this study. J.M. Martin-Garcia and P. Fromme designed all experiments of the cp-Taspase1_α41-233/β_ protein. J.M. Martin-Garcia, N. Nagaratnam, and R. Jernigan did the bioinformatics analyses. J.M. Martin-Garcia, N. Nagaratnam, and R. Jernigan crystallized and characterized crystals as well as collected data for the cp-Taspase1_α41-233/β_ protein. J.M. Martin-Garcia and D. Thifault processed anisotropic data of cp-Taspase1_α41-233/β_. J.M. Martin-Garcia and P. Fromme determined and evaluated the structure of cp-Taspase1_α41-233/β_ protein. D. Williams, B. Nannenga and N. Nagaratnam prepared and imaged samplesfor EM experiments. D. Williams analyzed negative staining EM experiments of cp-Taspase1_α41-233/β_. J.M. Martin-Garcia and N. Nagaratnam prepared all figures for the manuscript. J.M. Martin-Garcia, P. Fromme, N. Nagaratnam, and R. Jernigan wrote the manuscript with contributions from all co-authors.

