## Supplementary Material for "New Structural Insights into the Function of the Catalytically Active Human Taspase1"

#### Supplementary Information

##### Analysis of the interfaces observed in the crystal structure of the cp-Taspase1$_{\alpha41\text{-}233/\beta}$ protein

**<u>The dimer interface</u>:** The dimer interface within the αββα units is composed of 60 residues from chain A and 59 residues from chain B and buries a large surface area of 2039Å$^2$. This interface contains a complex network of 28 contacts (21 hydrogen bonds and 7 salt bridges) between the two monomers (Figure S18). As the core of the αββα unit is structurally very conserved between all current Taspase1 structures, this interface is very similar in all structures. All contacts are listed in Table S5.

**<u>The intra-ring and histidine interfaces</u>:** There are two type of interfaces that keep three αββα dimers together within each single-ring structure, which we have termed the 'intra-ring' interface, and the 'histidine' interface. Both interfaces are located between the β subunits within a single ring and bury a surface area of 1355Å$^2$ for the intra-ring interface and 69.7 Å$^2$ for the histidine interface (Table 2). At the intra-ring interface, we have identified two distinct patches: patch 1 that involves most residues at helix αH (between Ala312 and Lys326), and patch 2 that involves some of the residues at beta strands β10-β13 and the loops connecting them. Both patches are illustrated in Figure S19. The entire interface contains a total of 15 contacts (13 hydrogen bonds and 2 salt bridges, Table 2). The interface with all contacts is illustrated in Figure S20. All contacts are listed in Table S5. The other interface that maintains the single ring structure is the histidine interface (Figure S21). The two His281 residues located at the β subunit (one from each chain of the dimer) are pointing toward the center of the ring and interacting with Glu308 residues of β subunit from neighboring dimers as shown in Figure S21. There are 3 contacts found in this interface (Table S5).

**The double-ring interfaces:** Between opposite single-rings, the dimers interact at two interfaces that have been termed the ‘helix’ interface, and the ‘linker’ interface (Table 2). The helix interface comprises the long helical domain running from the lower ring and another one running from the upper ring (Figure S22). Due to the sequential up-and-down orientation of the helices in the single-ring, there are three helix interfaces per double-ring with 19 residues involved per helix and a buried surface area of 648.2$Å^2$. This interface is maintained through only two polar contacts (Figure S22 and Table S5). The other interface between opposite single-rings is the linker interface that comprises residues close to the GSGS linker (Figure S23). This interface has an accessible surface area of 168.0$Å^2$ and there are two inter-chain salt bridge contacts involved (Figure S23 and Table S5). Thus, the presence of only a few hydrogen bonds in the double-ring interface indicates that this interface is maintained mainly by hydrophobic and van del Walls interactions through the helices.

A visual inspection indicates loose packing of residues at each interface, and PDBePISA predicts favorable solvation energies between -1.7 kcal/mol and -23.8 kcal/mol for all of the interfaces (Table 2), and complex formation significance scores (CSS) that vary from 0 (for the helix, linker, histidine, and intra-ring interfaces) to 1 (for the dimer and intra-ring interfaces, Table 2). This indicates that weak hydrophobic interactions at the interfaces between single rings with low CSS (<0.1) very likely represent contacts formed during crystallization. In contrast, for those interfaces with high CSS of 1, the hydrophobic forces to keep interfaces together become significant, which indicates that both the dimer interface and the intra-ring interface are stable and might also exist in solution.

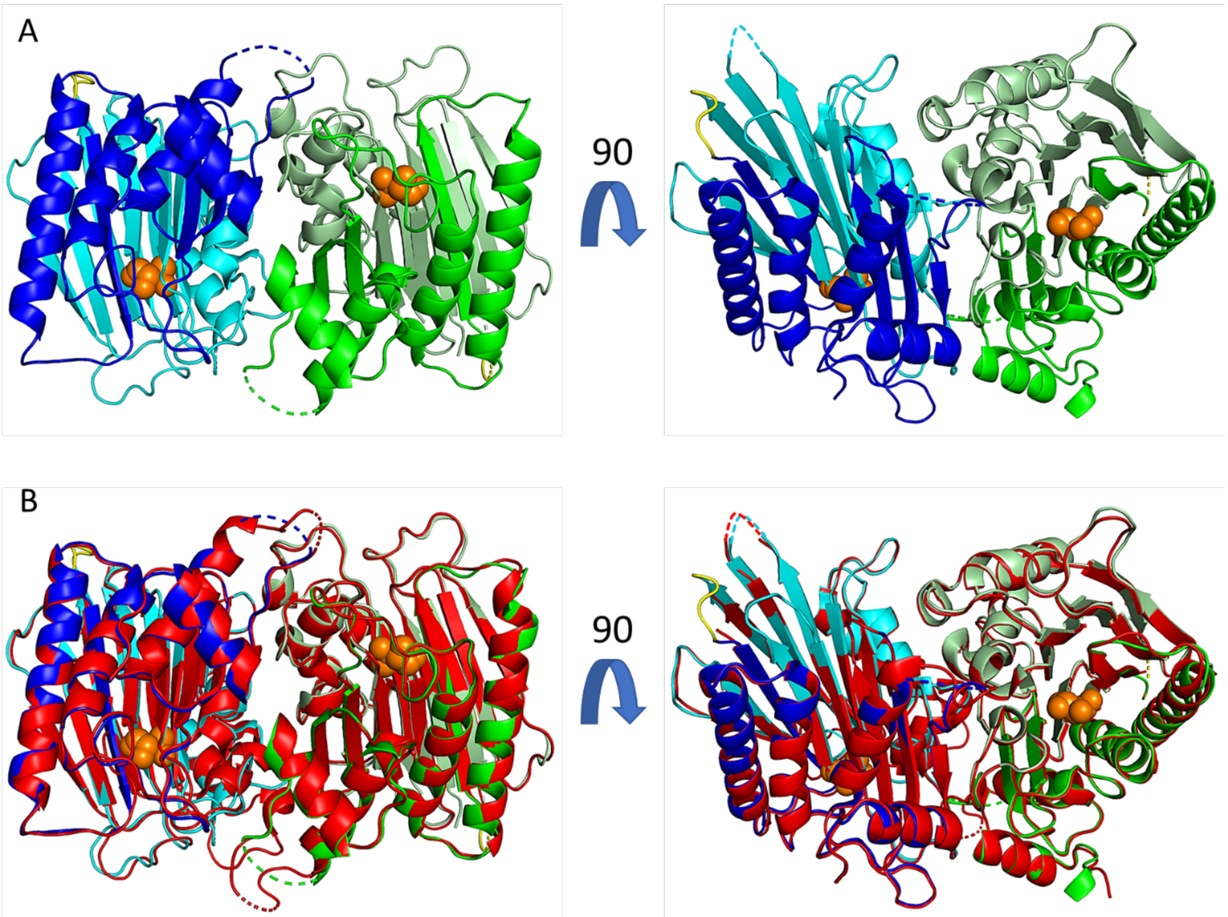


**Figure S1.** Crystal structure of cp-Taspase1$_{\alpha41\text{-}183/\beta}$. A) Side (left) and top (right) views of the hetero-tetramer in cp-Taspase1$_{\alpha41\text{-}183/\beta}$. Cartoon representation of the α subunit (green) and β subunit (light green) of chain A and α subunit (blue) and β subunit (cyan) chain B of the dimer. The nucleophile Thr234 is shown as orange spheres. The linker (GSGS) connecting the alpha and beta subunits is depicted in yellow. All missing segments are indicated by dashed lines. B) structural alignment of cp-Taspase1$_{\alpha41\text{-}183/\beta}$ with the truncated, 2-chain Taspase1 protein from Khan et al (PDB 2A8J). The cp-Taspase1$_{\alpha41\text{-}183/\beta}$ is shown as in A). The truncated 2-chain Taspase1 is show as red cartoon. All missing segments are indicated by dashed lines.

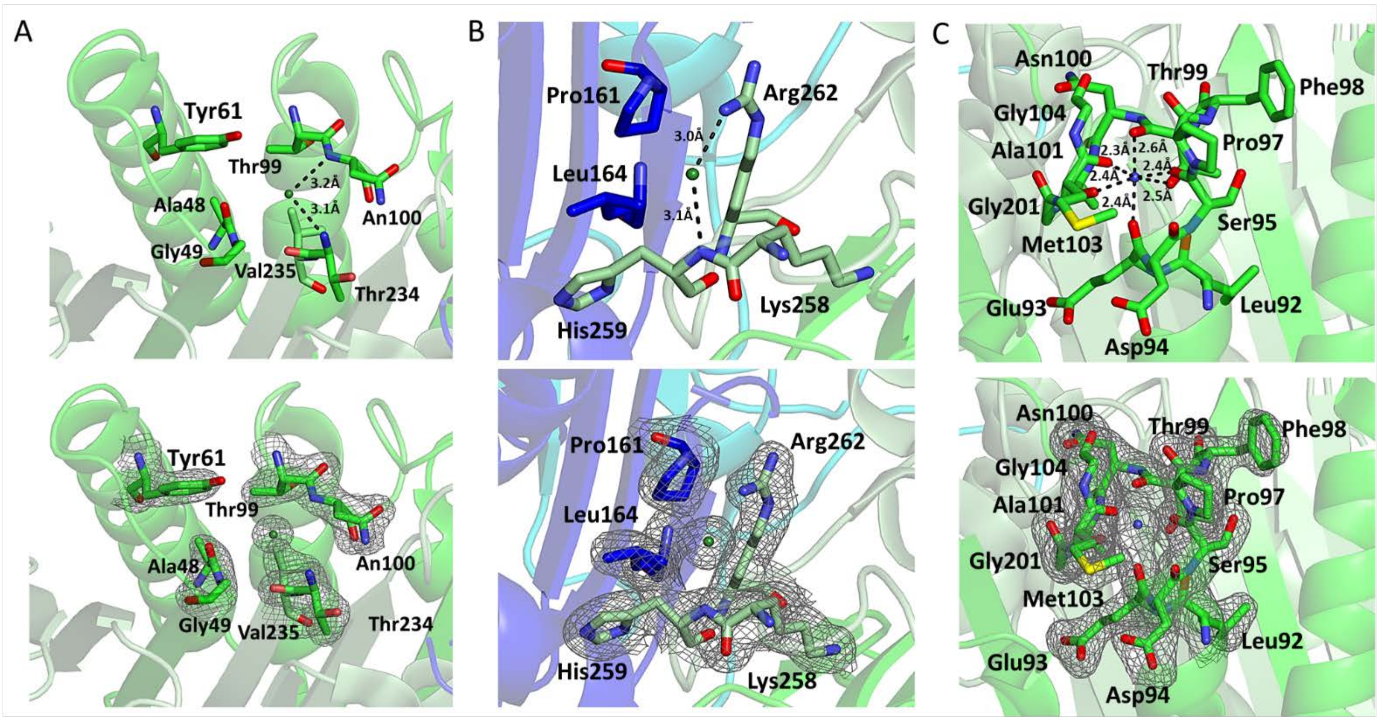


**Figure S2.** Ion binding sites in the crystal structure of cp-Taspase1$_{\alpha 41\text{-}183/\beta}$. A), B) Chloride ions, shown as green spheres, occupy the catalytic site (A) and a region adjacent to the active site (B). Interactions with protein residues are shown with black dashed lines. C) Sodium binding site. Sodium ion is shown as a blue sphere and its interactions with neighboring residues are indicated with black dashed lines. Distances to the ions have been shown. Lower panels show the electron density maps $2mF_o\text{-}DF_c$ contoured at $1\sigma$ around each binding site.

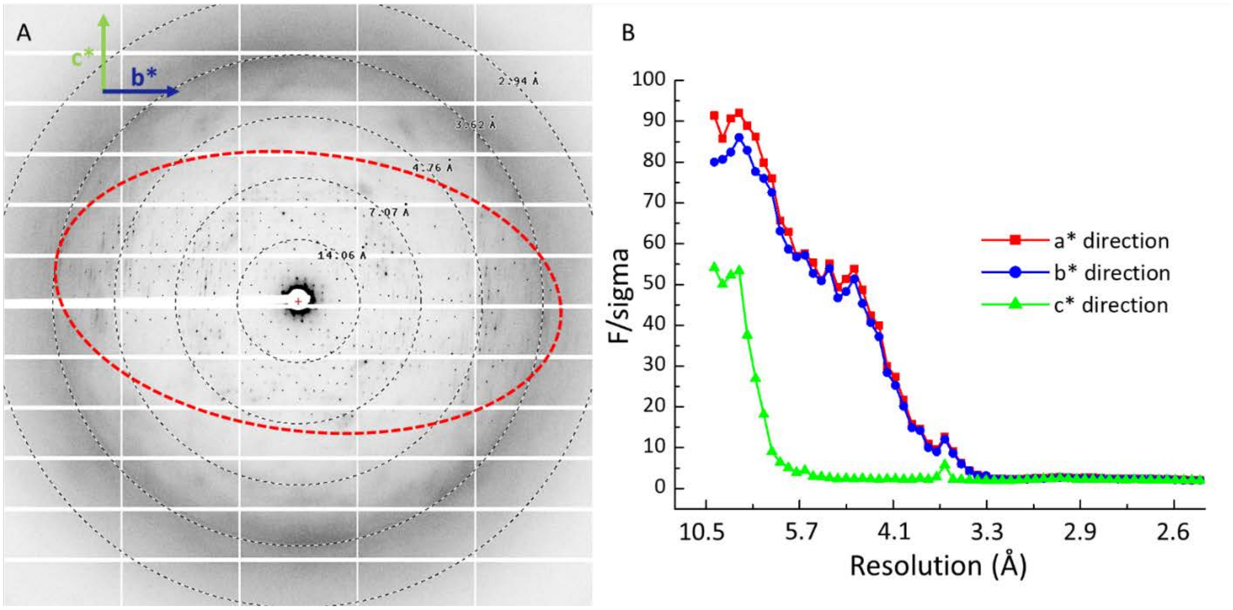


**Figure S3.** Anisotropy of cpTaspase1$_{\alpha41\text{-}233/\beta}$ crystals. A) Diffraction pattern showing the typical oval-shape indicative of anisotropy in cpTaspase1$_{\alpha41\text{-}233/\beta}$ crystals. Bragg reflections are seen to higher resolution in the horizontal direction than in the vertical direction. B) F/sigma vs resolution plot from TRUNCATE program (69) showing that cpTaspase1$_{\alpha41\text{-}233/\beta}$ crystals have two strong diffracting directions (a* and b*) and one very weak-diffracting direction (c*).

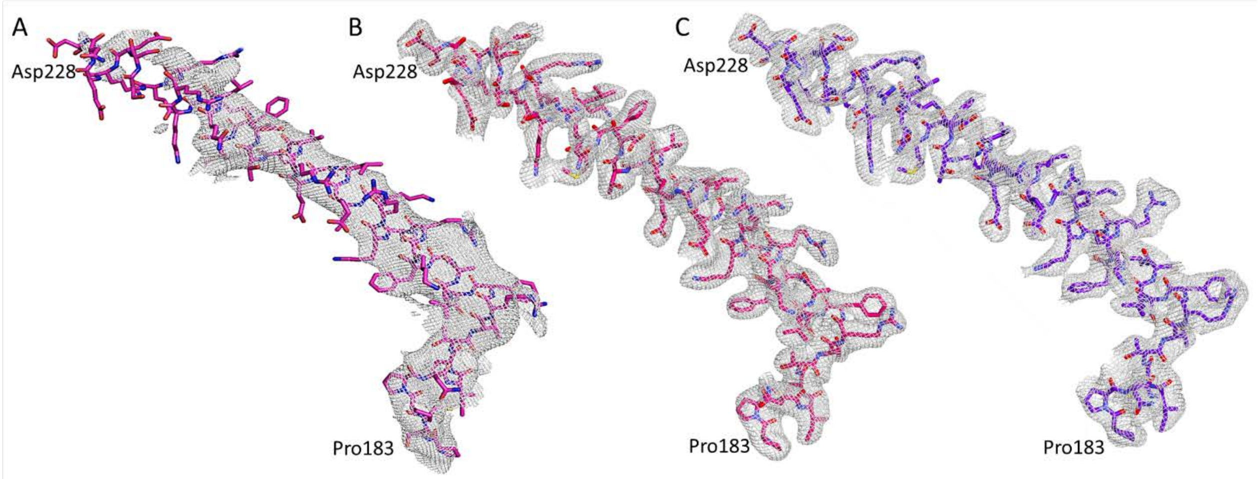


**Figure S4.** Electron density maps $2mF_o$-$DF_c$ contoured at $1\sigma$ of the long αE helix domain (residues Pro183-Asp228) of cpTaspase1$_{\alpha41\text{-}233/\beta}$. A) Electron density maps of chain A before anisotropy correction. B) Electron density map of chain A after anisotropy correction C) Electron density map of chain B after anisotropy correction.

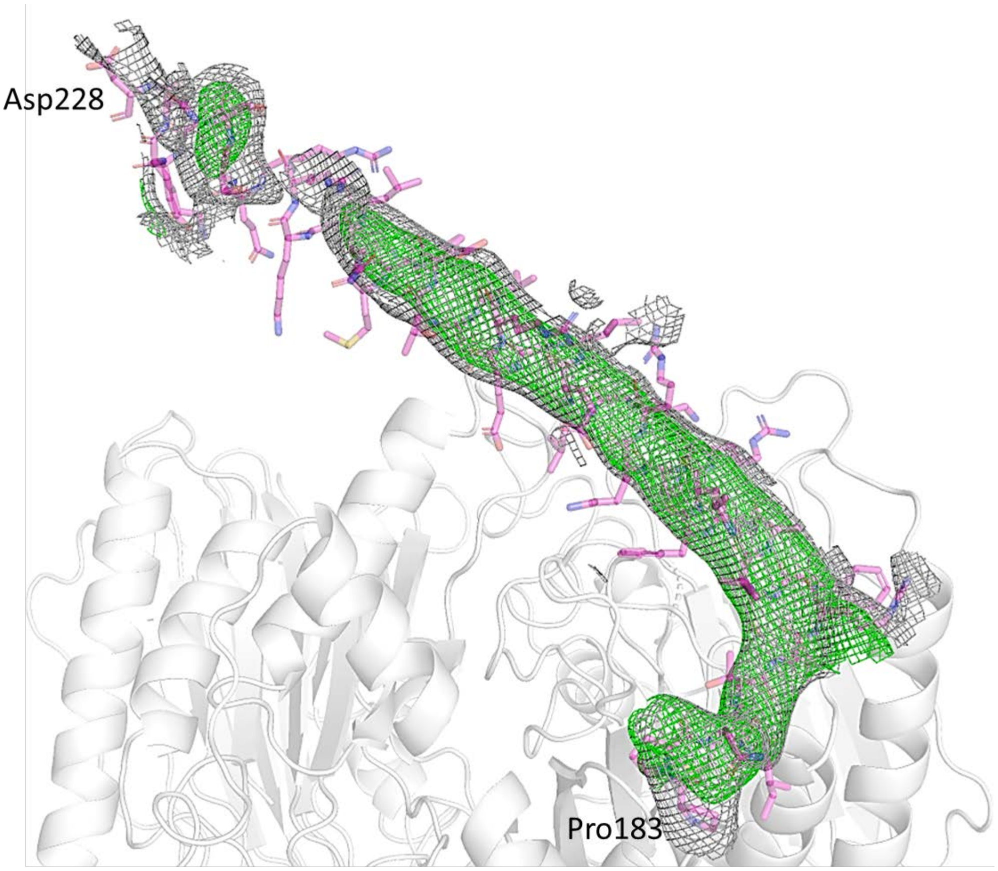


**Figure S5.** Electron density maps $2mF_o$-$DF_c$ (grey) and $mF_o$-$DF_c$ (green) contoured at 1σ and 3σ, respectively, of the long αE helix domain (residues Pro183-Asp228) of chain A of cpTaspase1$_{\alpha 41\text{-}233/\beta}$ after the first refinement step and before correcting anisotropy. The rest of the protein is shown as a white cartoon.

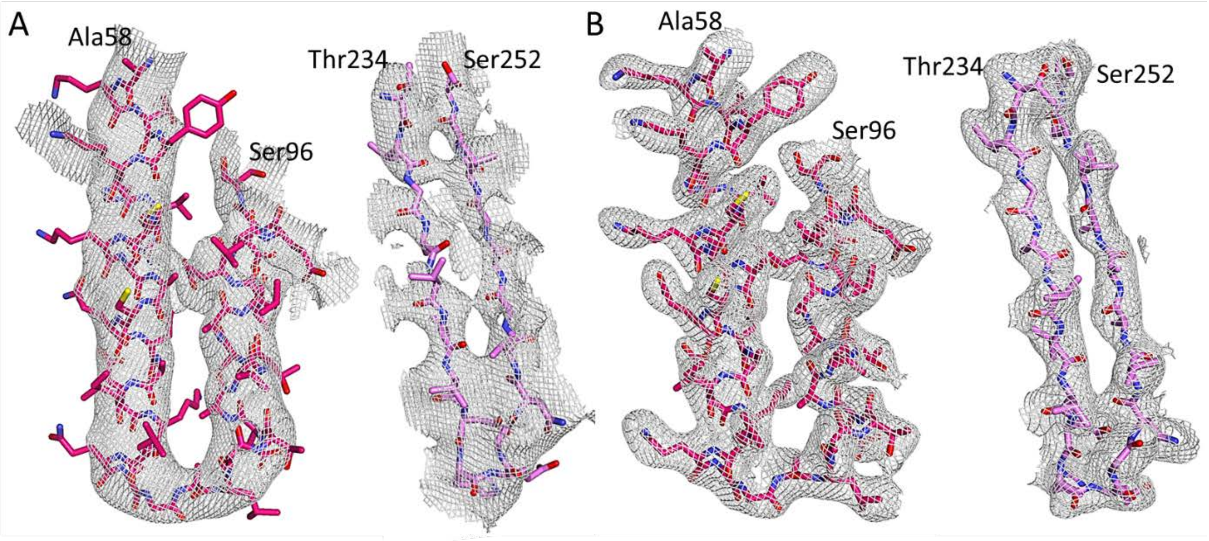


**Figure S6.** Electron density maps $2mF_o$-$DF_c$ contoured at 1σ of the regions Ala58-Ser96 (in the α-subunit) and Thr234-Ser22 (in the β-subunit) of chain A of cpTaspase1$_{\alpha 41\text{-}233/\beta}$ before (A) and after (B) anisotropy correction.

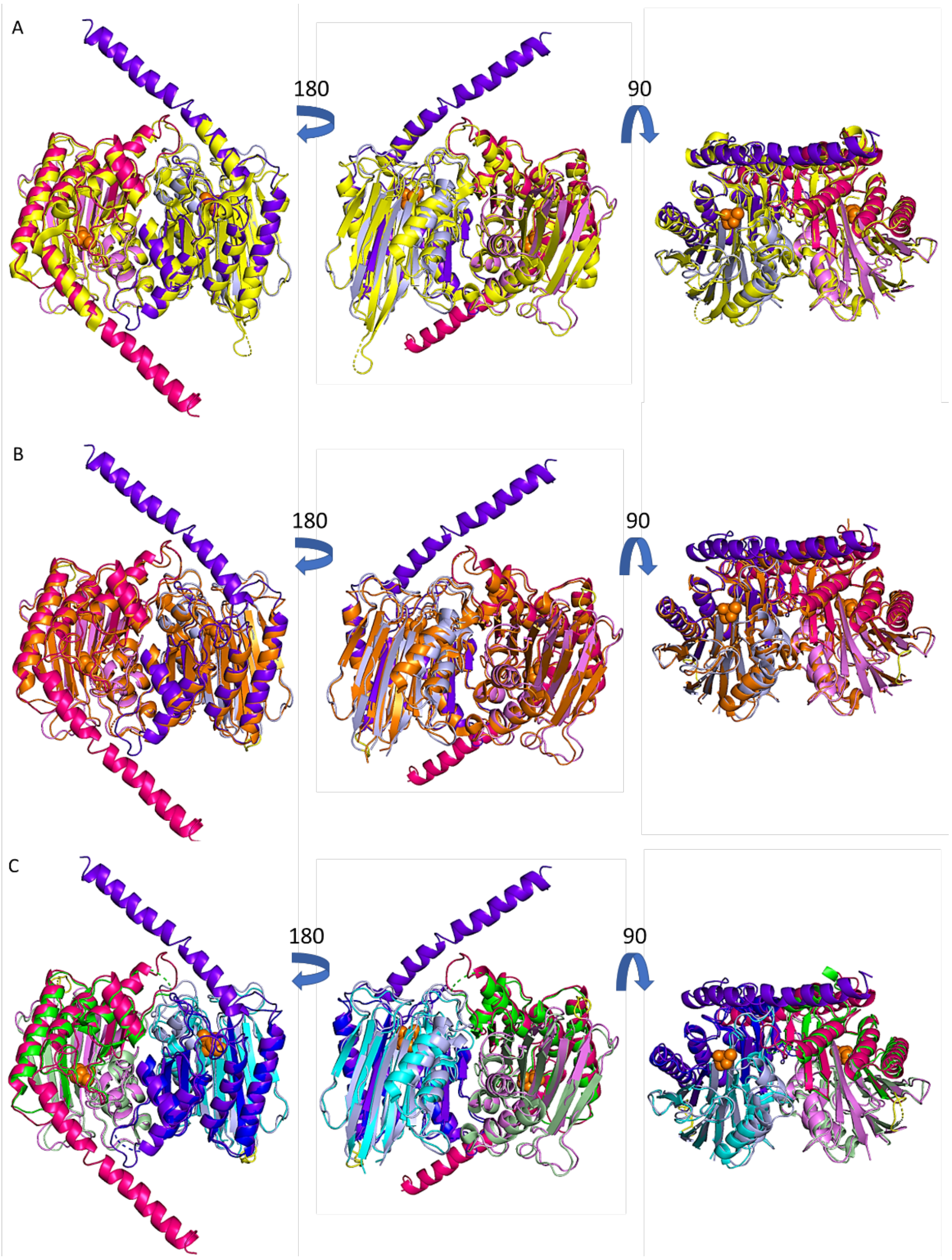


**Figure S7.** Structural comparison of the cpTaspase1$_{\alpha41\text{-}233/\beta}$ with other Taspase1 structures. A) Superposition of cpTaspase1$_{\alpha41\text{-}233/\beta}$ with the proenzyme (PDB 2A8I (14), B) Superposition of cpTaspase1$_{\alpha41\text{-}233/\beta}$ with truncated 2-chain Taspase1(PDB 2A8J (14), C) Superposition of cpTaspase1$_{\alpha41\text{-}233/\beta}$ with cpTaspase1$_{\alpha41\text{-}183/\beta}$. All structures are represented as cartoons using the following color code: cpTaspase1$_{\alpha41\text{-}233/\beta}$ (pink (α-subunit), and magenta (β-subunit) for chain A, and purple (α-subunit), and light blue (β-subunit) for chain B), cpTaspase1$_{\alpha41\text{-}183/\beta}$ (green(α-subunit), and light green (β-subunit) for chain A, and blue (α-subunit), and cyan (β-subunit) for chain B), proenzyme (yellow), and truncated Taspase1 (orange). The nucleophile Thr234 is represented as with orange spheres.

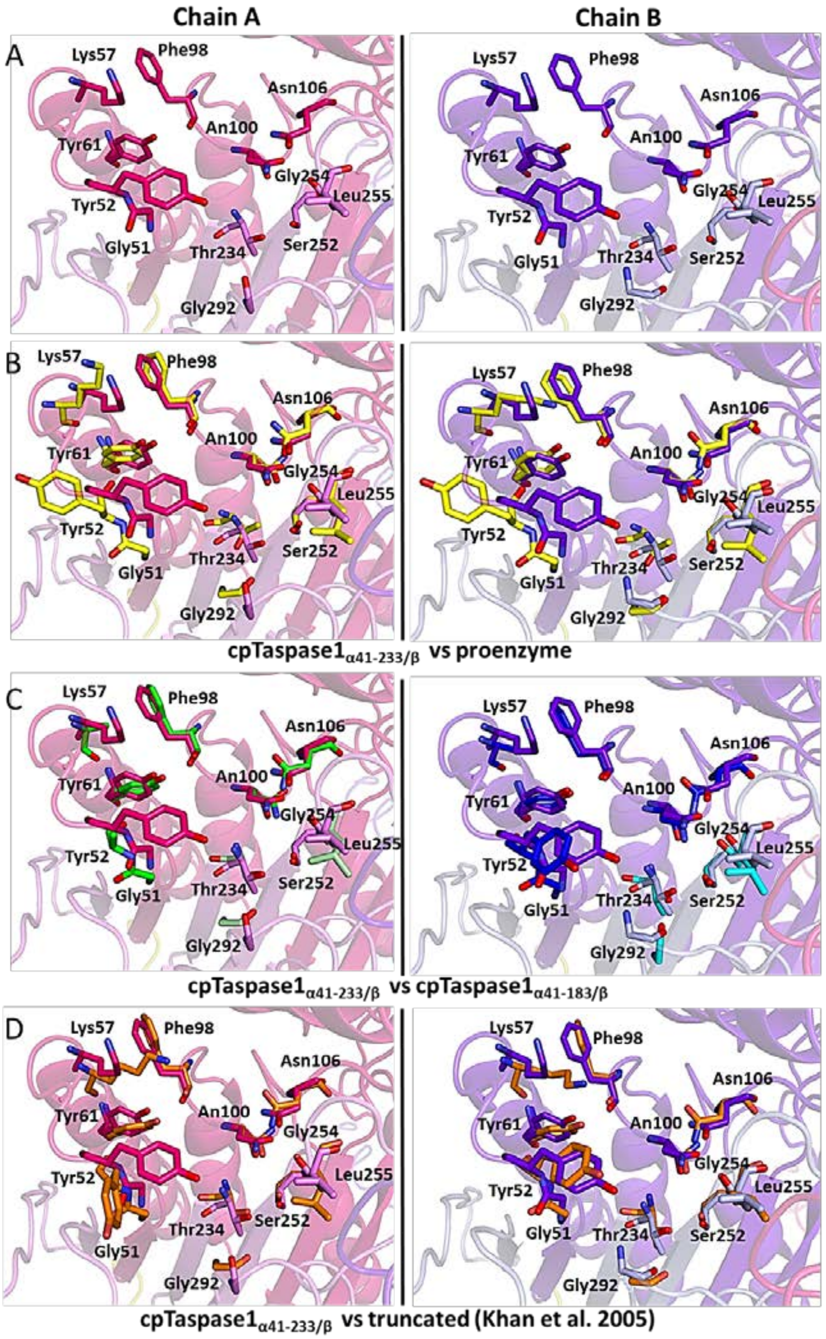


**Figure S8.** Catalytic site of chains A and B of cpTaspase1$_{\alpha41\text{-}233/\beta}$ and their comparison to other Taspase1 structures. A) Residues at the catalytic site of chains A and B of cpTaspase1$_{\alpha41\text{-}233/\beta}$ alone. B) Superimposition of chains A and B (as shown in panel A) with the proenzyme (PDB 2A8I (14). C) Superimposition of chains A and B (as shown in panel A) to cpTaspase1$_{\alpha41\text{-}183/\beta}$. D) Superimposition of chains A and B (as shown in panel A) with the truncated Taspase1 (PDB 2A8J (14)).

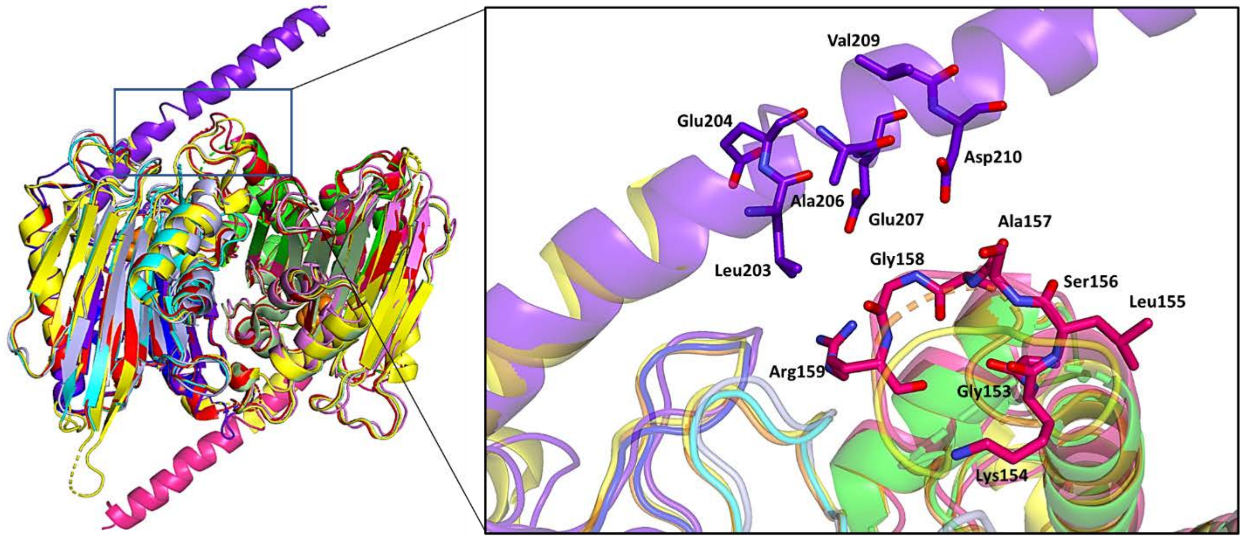


**Figure S9.** Structural differences at loop Gly153-Arg159. Left: Superposition of cpTaspase1$_{\alpha41\text{-}233/\beta}$ with cpTaspase1$_{\alpha41\text{-}183/\beta}$, the proenzyme (PDB 2A8I (14), and truncated 2-chain Taspase1(PDB 2A8J (14). Right: Residues of the fragment Gly153-Arg159 fully modeled in cpTaspase1$_{\alpha41\text{-}233/\beta}$ compared to that in other Taspase1 structures. Dashed lines indicate missing residues in the loop. Neighboring residues in the middle of the long domain are also shown. All structures are represented as cartoons using the same color code as in Figure S8.

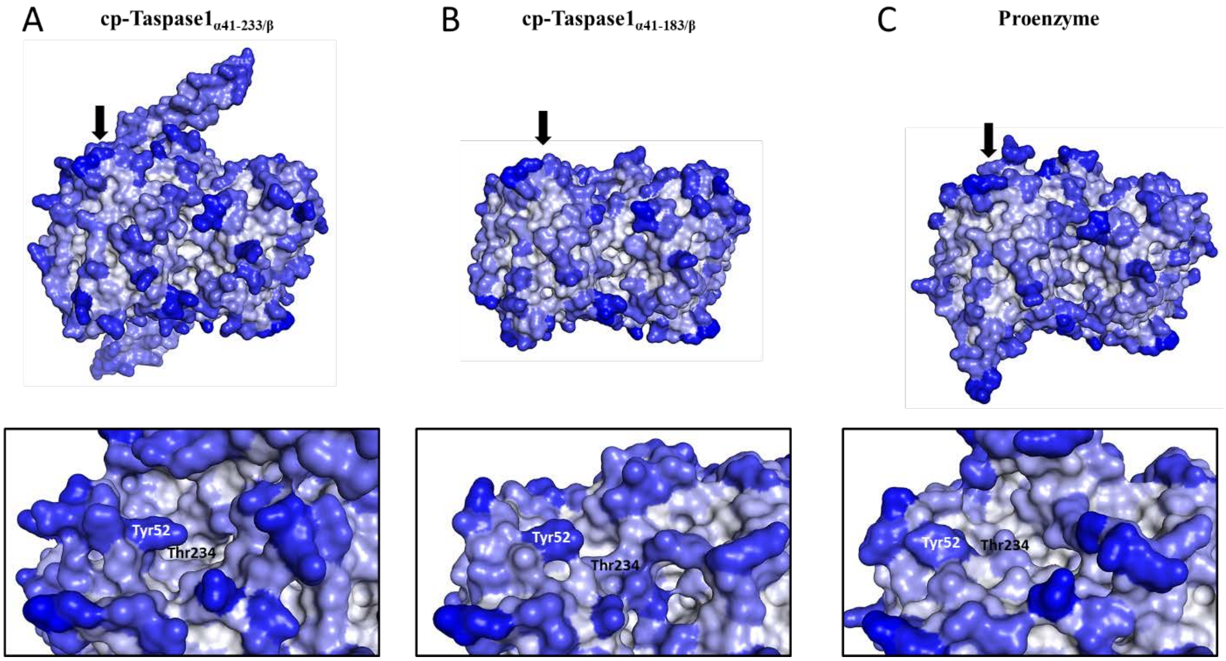


**Figure S10.** Accessible surface area (ASA) of cp-Taspase1$_{\alpha 41\text{-}233/\beta}$ (A), cp-Taspase1$_{\alpha 41\text{-}183/\beta}$ (B), and the proenzyme (PDB 2A8I(14)) (C). ASAs per residue are illustrated using a scale from 0 (white) to 1 (dark blue). Residues with low ASA values are less accessible than those with high ASA values. Upper panels represent an overview of the structures and the catalytic sites are indicated by the arrows. Lower panels represent a closer-up view of the catalytic sites. The nucleophile Thr234 and the residue Tyr52 are shown as references.

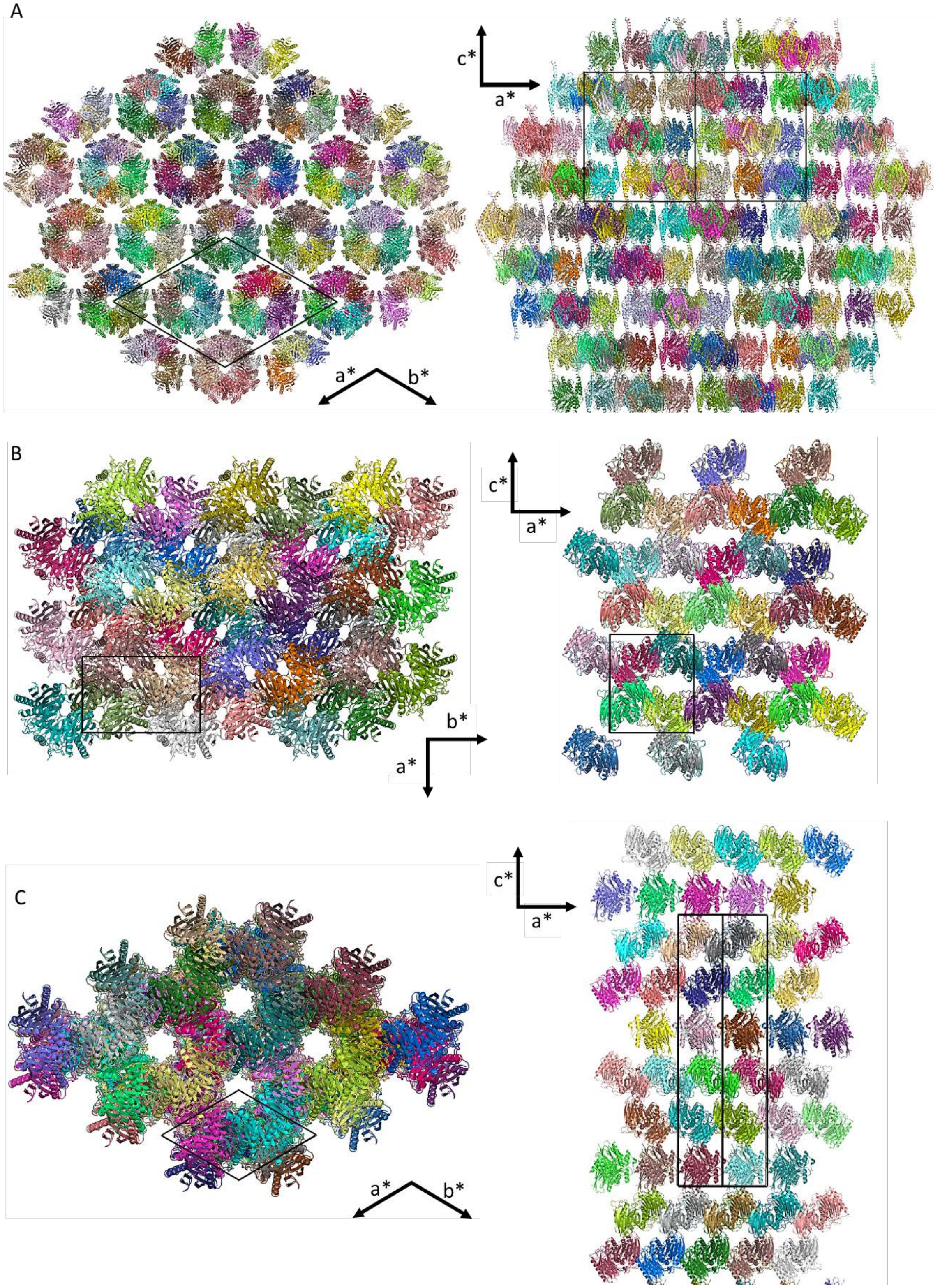


**Figure S11.** Orthographic projection of the crystal packing represented in two different directions for the double-ring assembly of cpTaspase1$_{\alpha 41\text{-}233/\beta}$, (A), the truncated 2-chain Taspase1(PDB 2A8J (14) (B), and the cpTaspase1$_{\alpha 41\text{-}183/\beta}$ (C). A unit cell is outlined for each view with black lines.

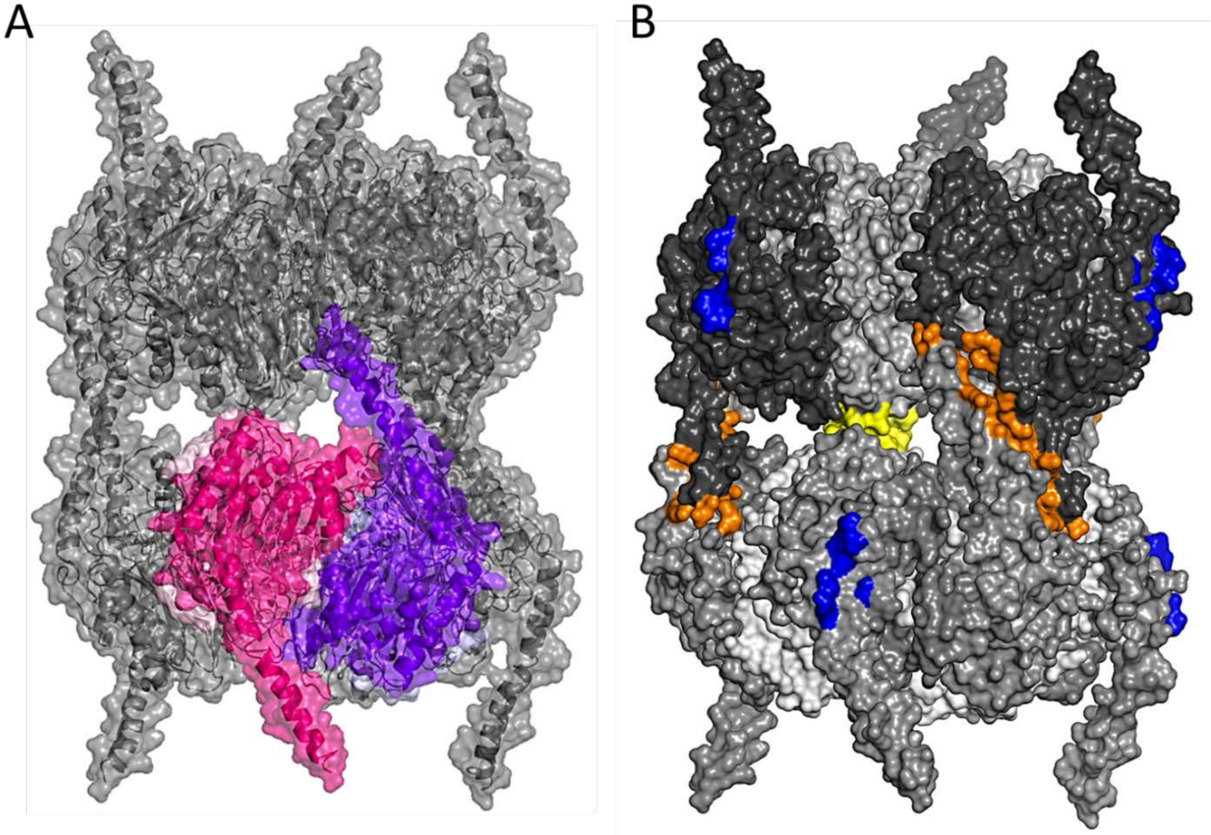


**Figure S12.** Subunit interfaces that maintain the double-ring assembly of cpTaspase1$_{\alpha41\text{-}233/\beta}$ in the crystals. A) Side view of the surface representation of the cpTaspase1$_{\alpha41\text{-}233/\beta}$ of the double-ring structure. B) Interfaces at the double-ring structure. The interfaces are colored as follows: orange for the helix interface, yellow for the linker interface, and blue for the inter-ring interface. The cpTaspase1$_{\alpha41\text{-}233/\beta}$ dimers are represented with the same color as in Figure 3 and the dimers generated by symmetry are shown in a grey and black.

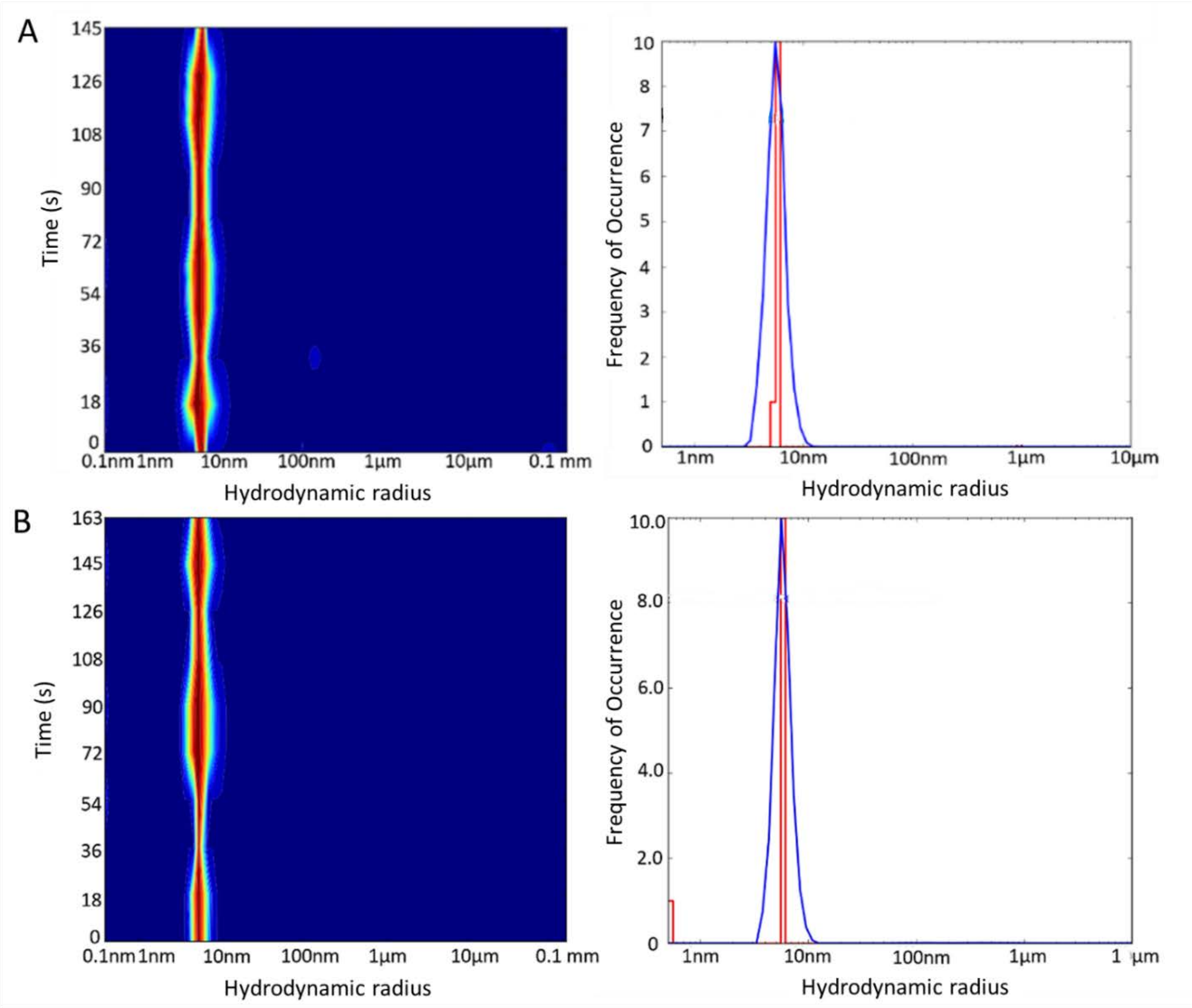


**Figure S13.** Dynamic light scattering (DLS) of cpTaspase1$_{\alpha 41\text{-}233/\beta}$. Size distribution (left panels) and histogram (right panels) plots show a very narrow size distribution. A) Experiment at 5 mg/ml. Hydrodynamic radius was 5.3 nm, which is equivalent to a molecular weight of ~183 kDa B) Experiment at 10 mg/ml. Hydrodynamic radius was 5.6 nm, which is equivalent to a molecular weight of ~220 kDa. The results shown are the average of ten measurements of 20 sec each. Experiment conditions were 20 mM HEPES pH 7.5, 5% glycerol, 300 mM NaCl.

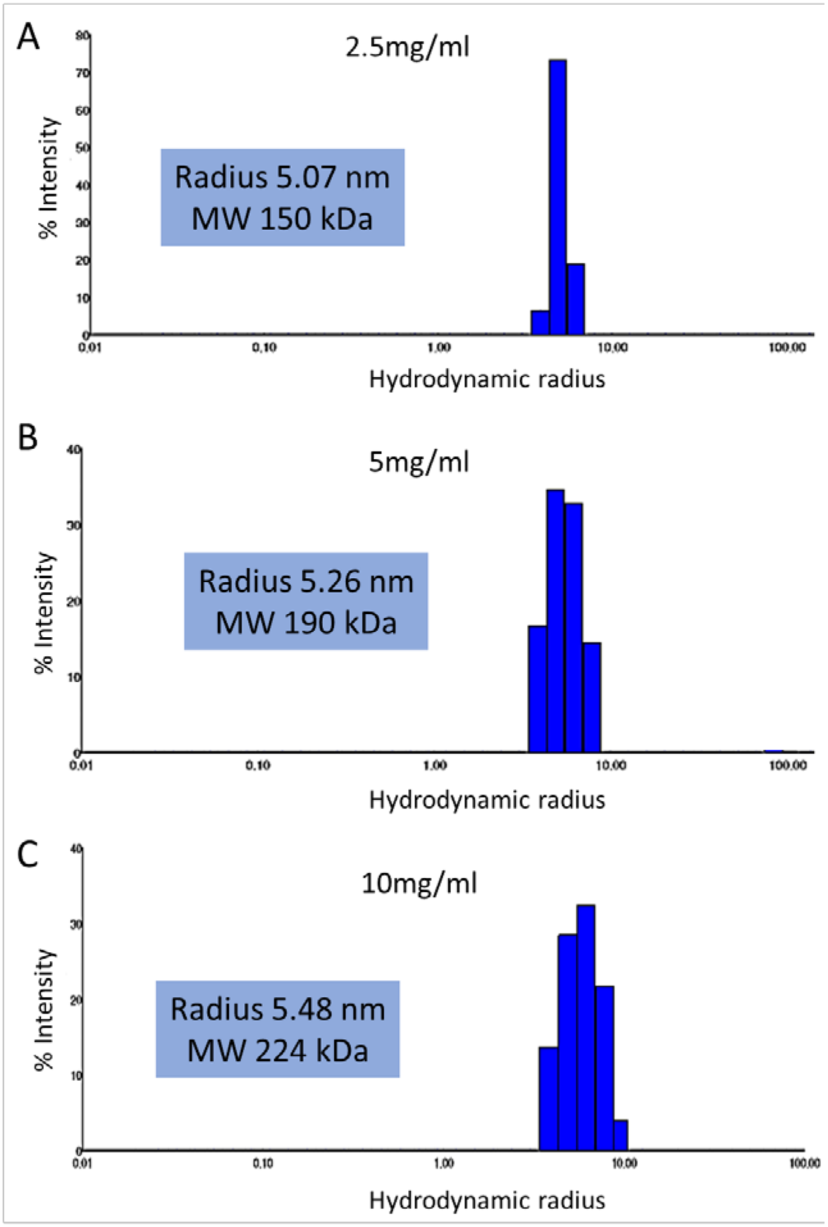


**Figure S14.** Dynamic light scattering (DLS) of cpTaspase1$_{\alpha 41\text{-}233/\beta}$. Histogram plots show a very narrow size distribution. A) Experiment at 2.5 mg/ml. Hydrodynamic radius was 5.07 nm, which is equivalent to a molecular weight of 150 kDa. B) Experiment at 5 mg/ml. Hydrodynamic radius was 5.26 nm, which is equivalent to a molecular weight of 190 kDa C) Experiment at 10 mg/ml. Hydrodynamic radius was 5.48 nm, which is equivalent to a molecular weight of 224 kDa. The results shown are the average of ten measurements of 20 sec each. Experiment conditions were 20 mM HEPES pH 7.5, 5% glycerol, 300 mM NaCl.

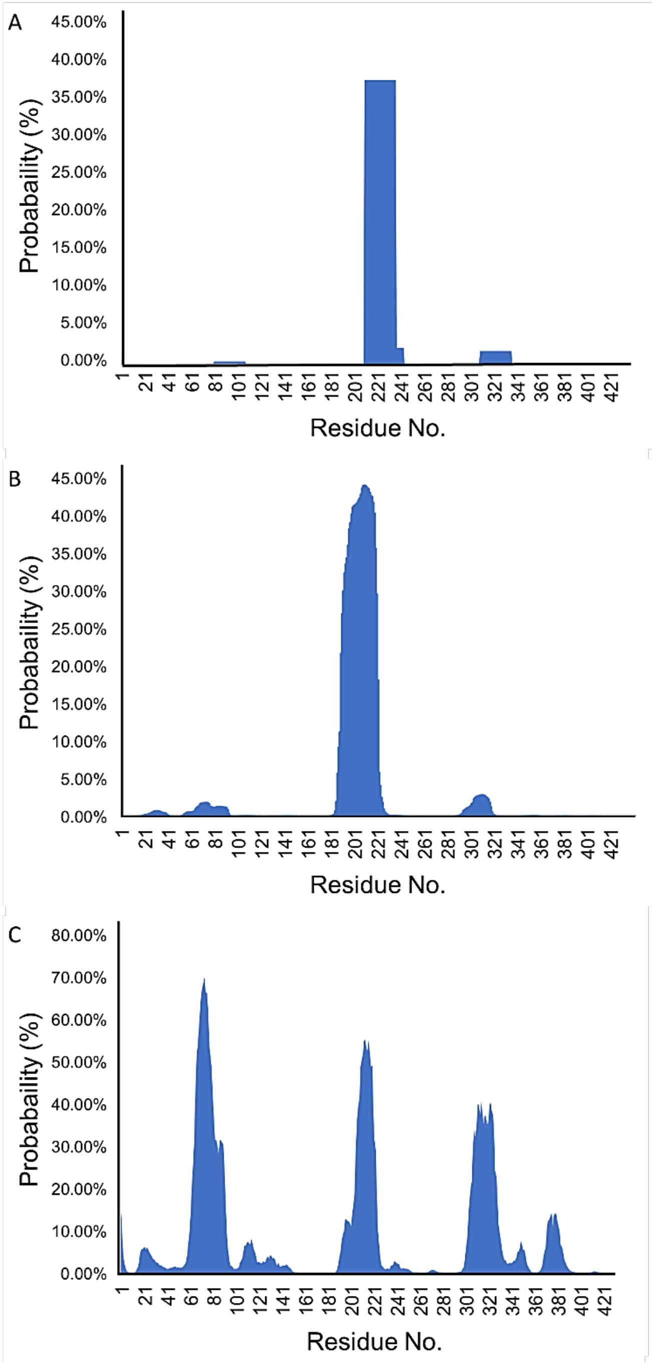


**Figure S15.** Coiled coil predictions for WT Taspase1 using COILS (25), MARCOIL (27) and DEEPCOIL (26) programs.

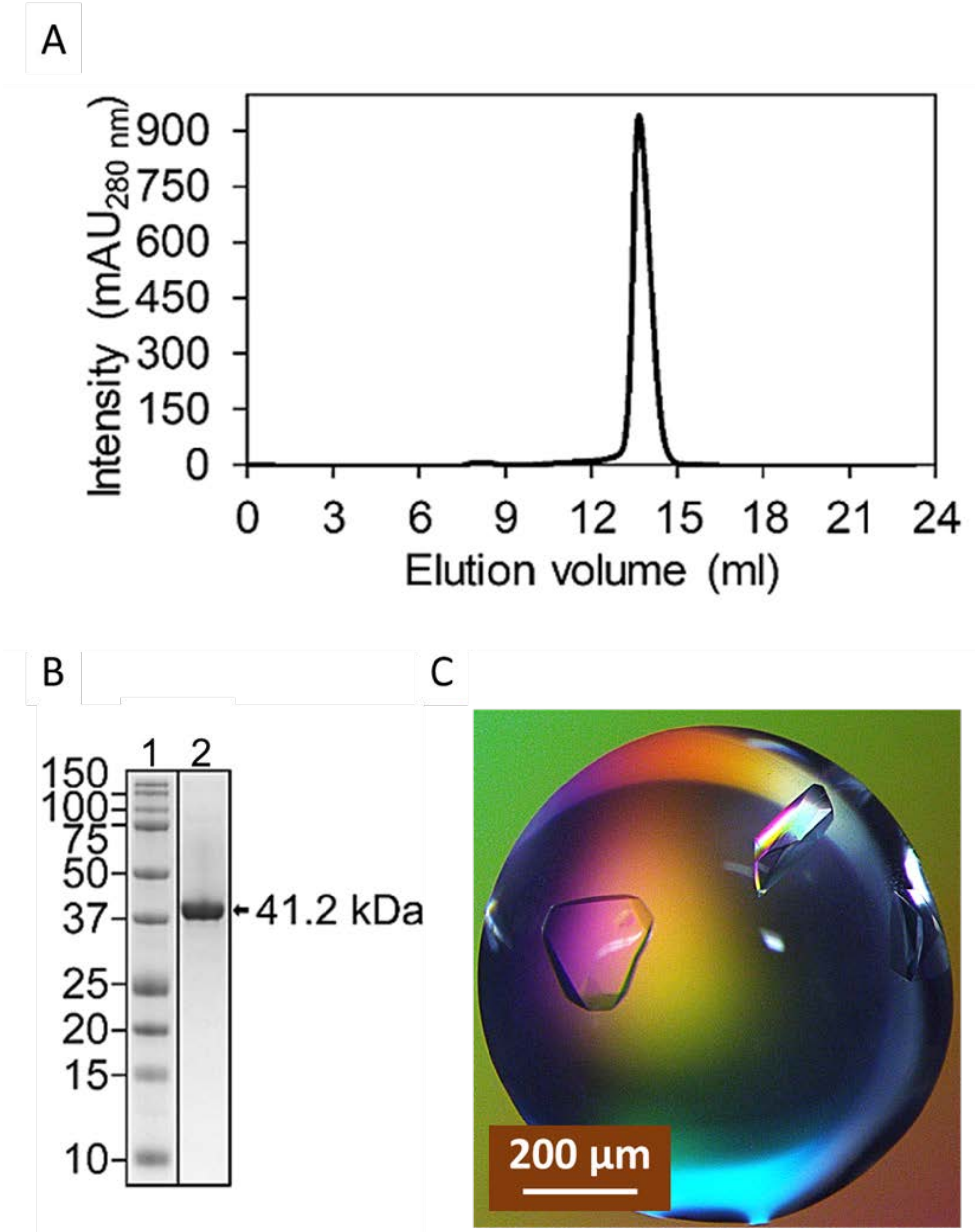


**Figure S16.** Purification and crystallization of the circularly permuted Taspase1, cpTaspase1$_{\alpha41\text{-}233/\beta}$. A) Size exclusion chromatography profile. The cpTaspase1$_{\alpha41\text{-}233/\beta}$ is eluted as a single peak at 13.6 ml elution volume, which suggests that the protein is highly monodisperse. B) SDS-PAGE analysis of eluted cpTaspase1$_{\alpha41\text{-}233/\beta}$ (lane 2). Observation of a single band between 37 and 50 kDa confirms the high purity of cpTaspase1$_{\alpha41\text{-}233/\beta}$ protein. Molecular weight markers are shown in lane 1. C) Crystals of cpTaspase1$_{\alpha41\text{-}233/\beta}$ grown in 0.1 M sodium citrate pH 4.0, 1.0 M ammonium sulfate, 10% 2-methyl-2,4-pentanediol (MPD), and 0.4 M NaCl, using the hanging drop vapor diffusion technique.

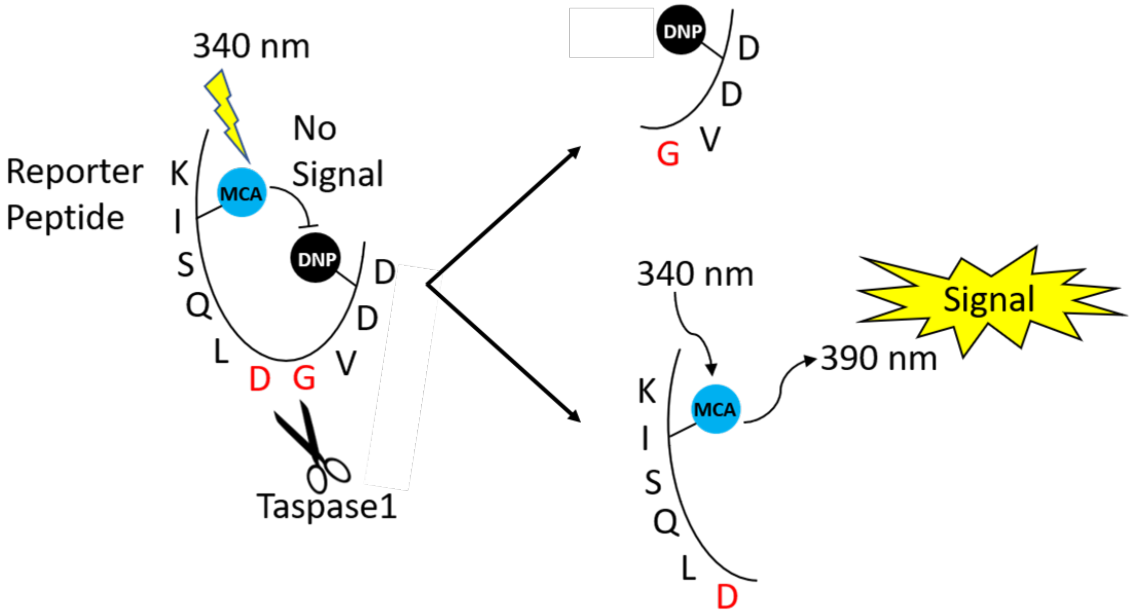


**Figure S17.** Schematic representation of the proteolytic cleavage reaction of the labeled peptidic substrate MCA-Lys-Ile-Ser-Gln-Leu-Asp↓Gly-Val-Asp-Asp-Lys(DNP)-NH2 by Taspase1. MCA and DNP are the fluorophore and the quencher, respectively. In the absence of Taspase1, the fluorophore does not emit due to close proximity of the quencher. When Taspase1 cleaves the peptide, the quencher is released from the fluorophore allowing emission of a fluorescent signal. Fluorescence intensities were measured with excitation 340/emission 455 filters using a Perkin Elmer Envision multi-mode plate reader and 384-well standard depth MCA_DNP protocol. Figure adapted from (62).

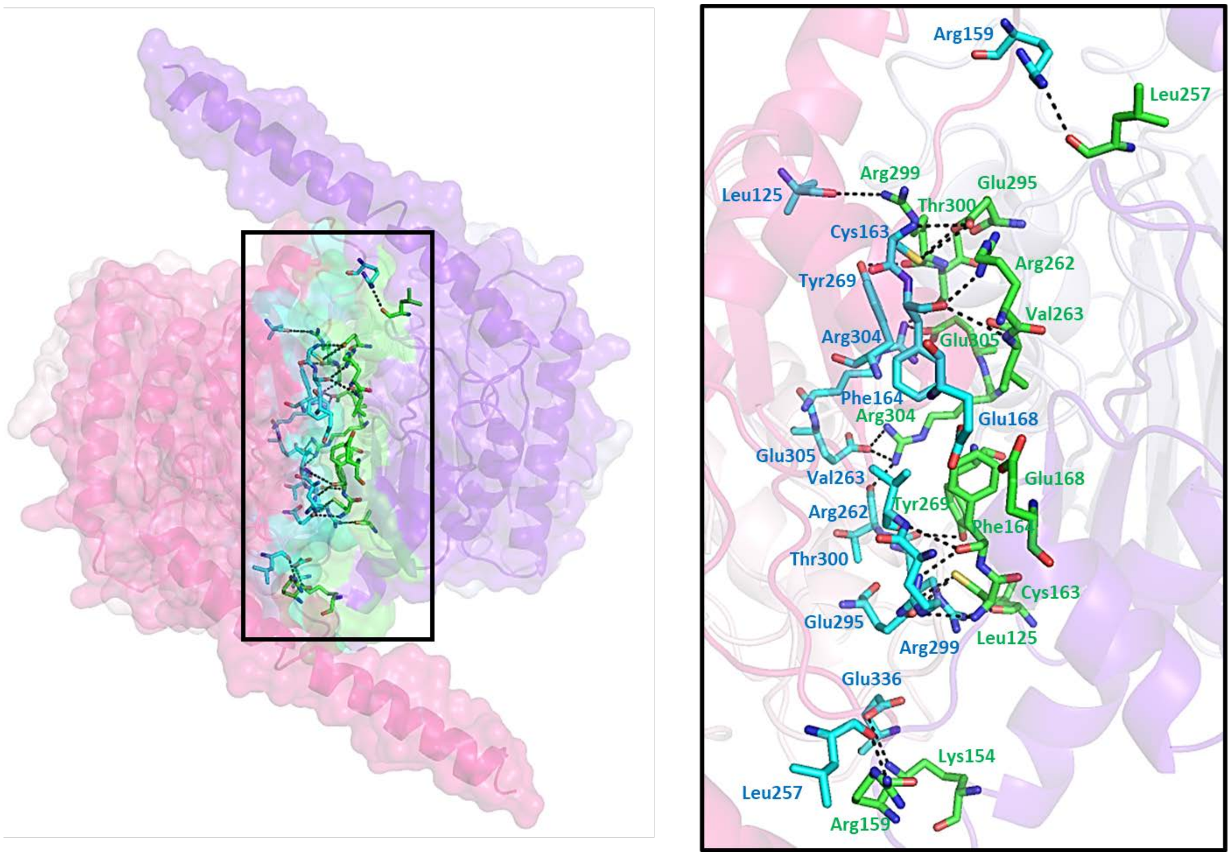


**Figure S18.** Dimer interface of cpTaspase1$_{\alpha41\text{-}233/\beta}$. Left: Cartoon and surface representation of the dimer in the asymmetric unit. The interface is highlighted in cyan and green. Right: Close-up view of the interface in the box on the left. Only residues involved in contacts are shown. All contact distances are shown as black dashed lines, and the numerical values are provided in Table S12.

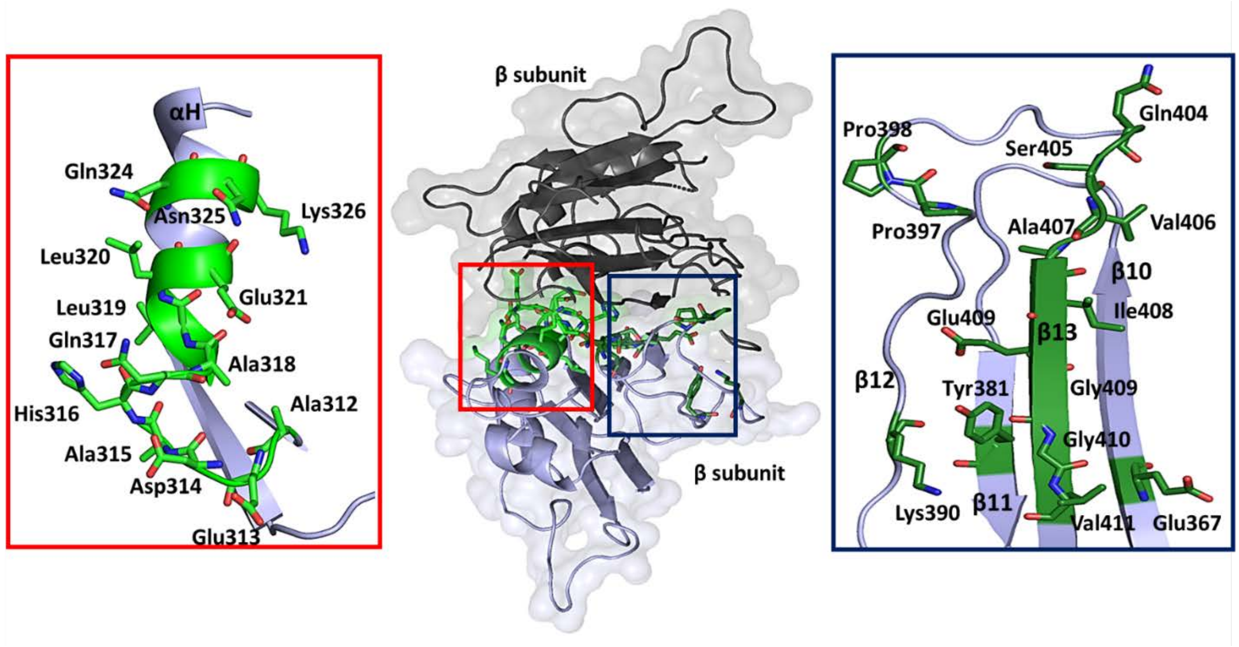


**Figure S19.** Inter-ring interface of cpTaspase1$_{\alpha41\text{-}233/\beta}$. Middle shows an overview of the interface between the two β-subunits highlighted light blue and black. The two patches found at the interface are in red and blue boxes. Left panel shows an expanded view of patch 1 highlighted green and shown as cartoon representation. Right panel shows a closer view of patch 2 highlighted dark green and shown as cartoon representation. For both patches, all residues involved in the interface are highlighted and shown in stick representation.

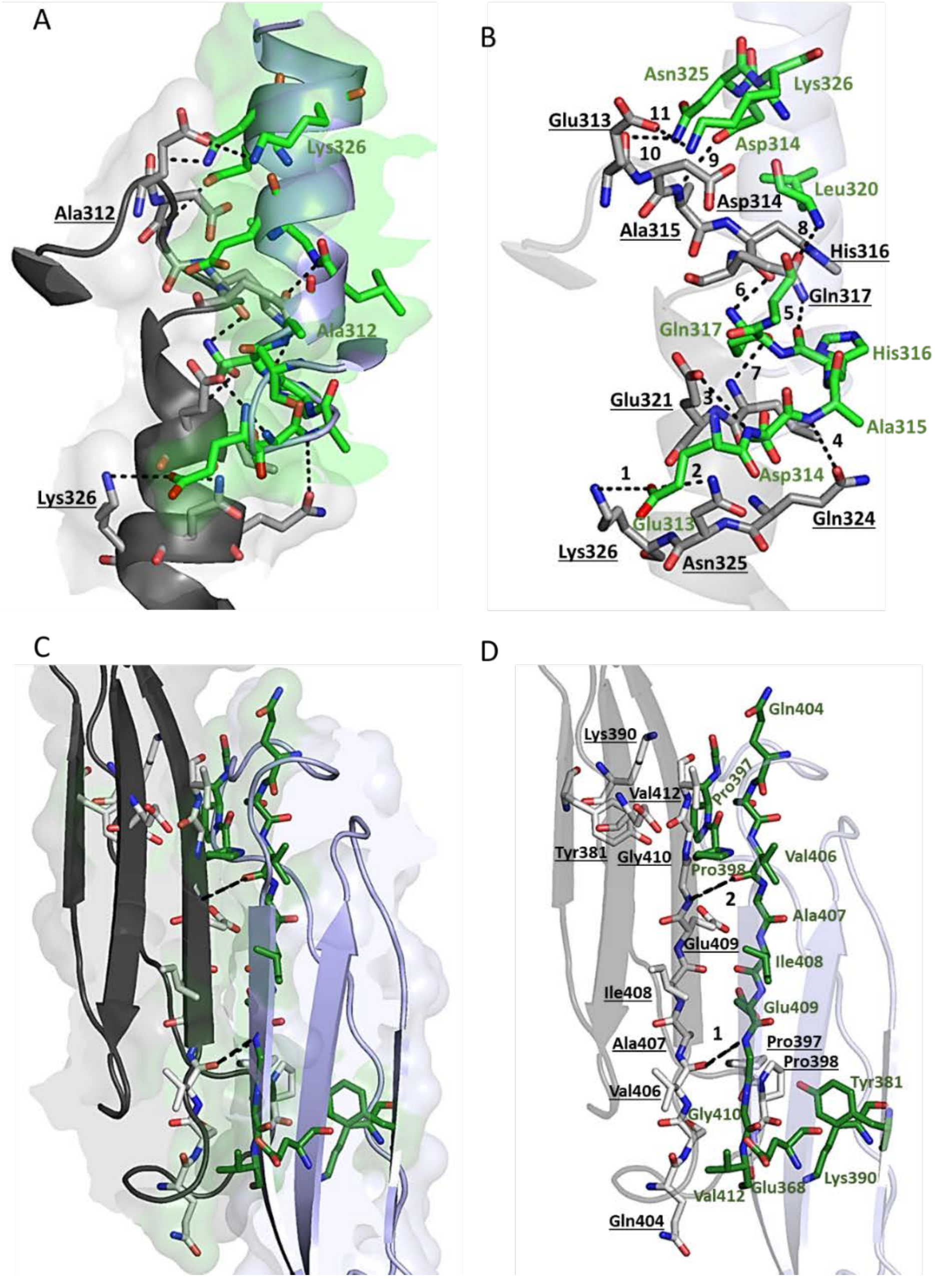


**Figure S20.** Crystal contacts at patches 1and 2 of the intra-ring interface of cpTaspase1$_{\alpha41\text{-}233/\beta}$. A) Overview representation of patch 1. The symmetry related fragment is shown in black. Residues at the interfaces are shown as sticks in green or grey (for the symmetry related fragment), and contacts as black dashed lines. B) Same representation as A). For clarity, residues not involved in any contact have been deleted. Contact distances are shown as black dashed lines and numbered according to that in Table S12. C), D) Overview representation of patch 2. Surface representation of the interface is shown in C. The symmetry related fragment is shown in black. Residues involved in the patch are represented as sticks in dark green and white (for the symmetry related fragment). The contacts at the interface are highlighted, shown as black dashed lines and numbered according to that in Table S12. For clarity, residue names and numbers from the symmetry related molecule are underlined.

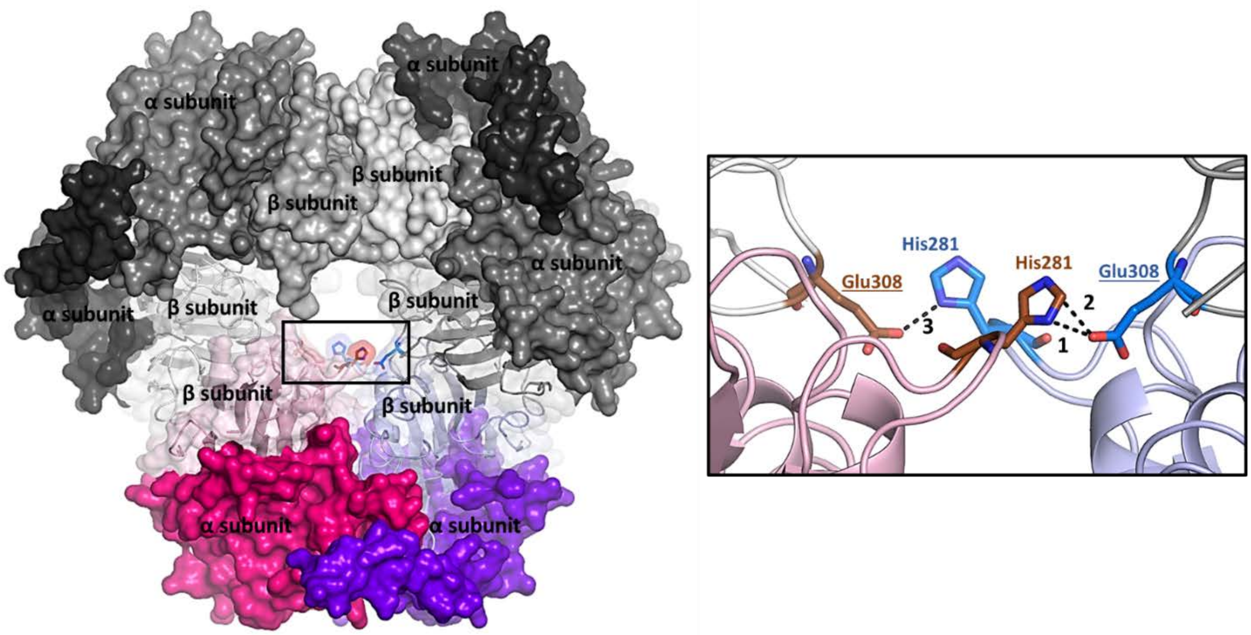


**Figure S21.** Histidine interface of cpTaspase1$_{\alpha41\text{-}233/\beta}$. Left: Top view of the single-ring structure with the histidine interface in black box. The dimer of the asymmetric unit is highlighted in color. The symmetry-related dimers are shown in black and grey. The α and β subunits are also labeled. Right: Detailed view of the contacts at the histidine interface. Residues at the interface are highlighted in red and blue. Contact distances are shown as black dashed lines and numbered according to that in Table S12. For clarity, Glu308 residues from the symmetry-related molecules are underlined.

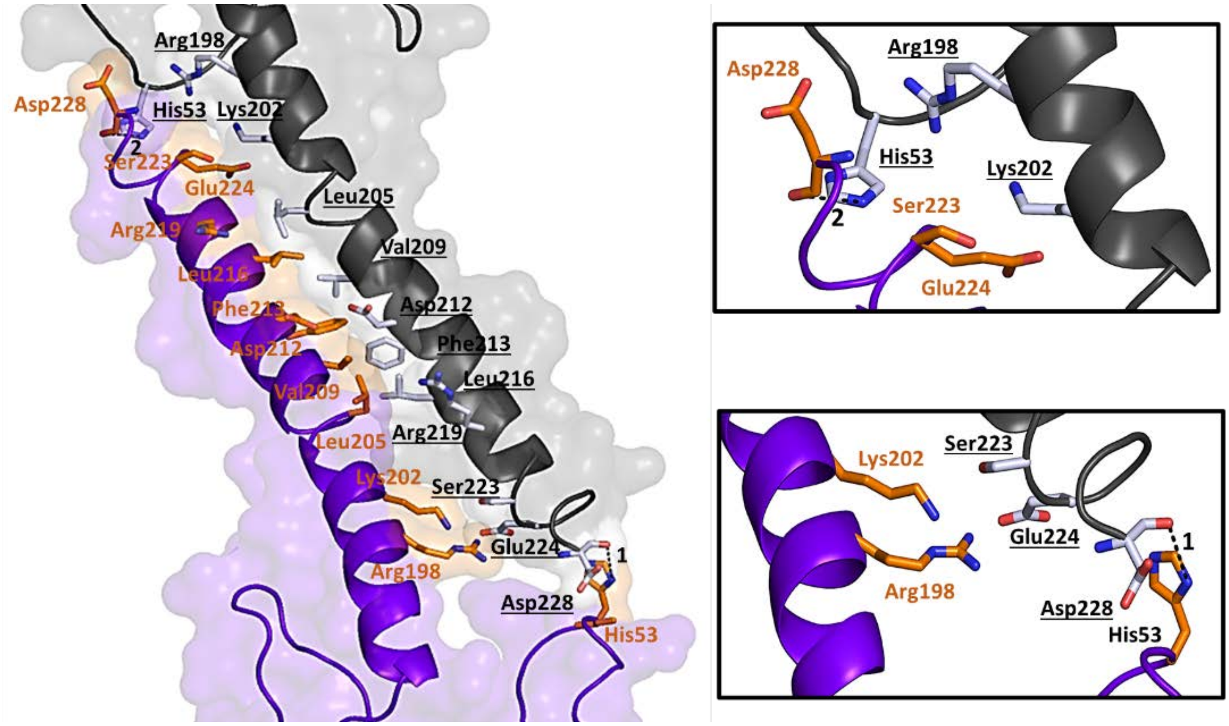


**Figure S22.** Helix interface of cpTaspase1$_{\alpha 41\text{-}233/\beta}$. Cartoon and surface representation of the long αE helices running from the lower ring (purple) and the upper ring (black). Residues involved in the interface are highlighted in orange and white and represented as sticks. The two contacts are also highlighted and represented as back dashed lines. Boxed panels show a close-up view of the contacts. For clarity, residue names and numbers from the symmetry related molecule are underlined.

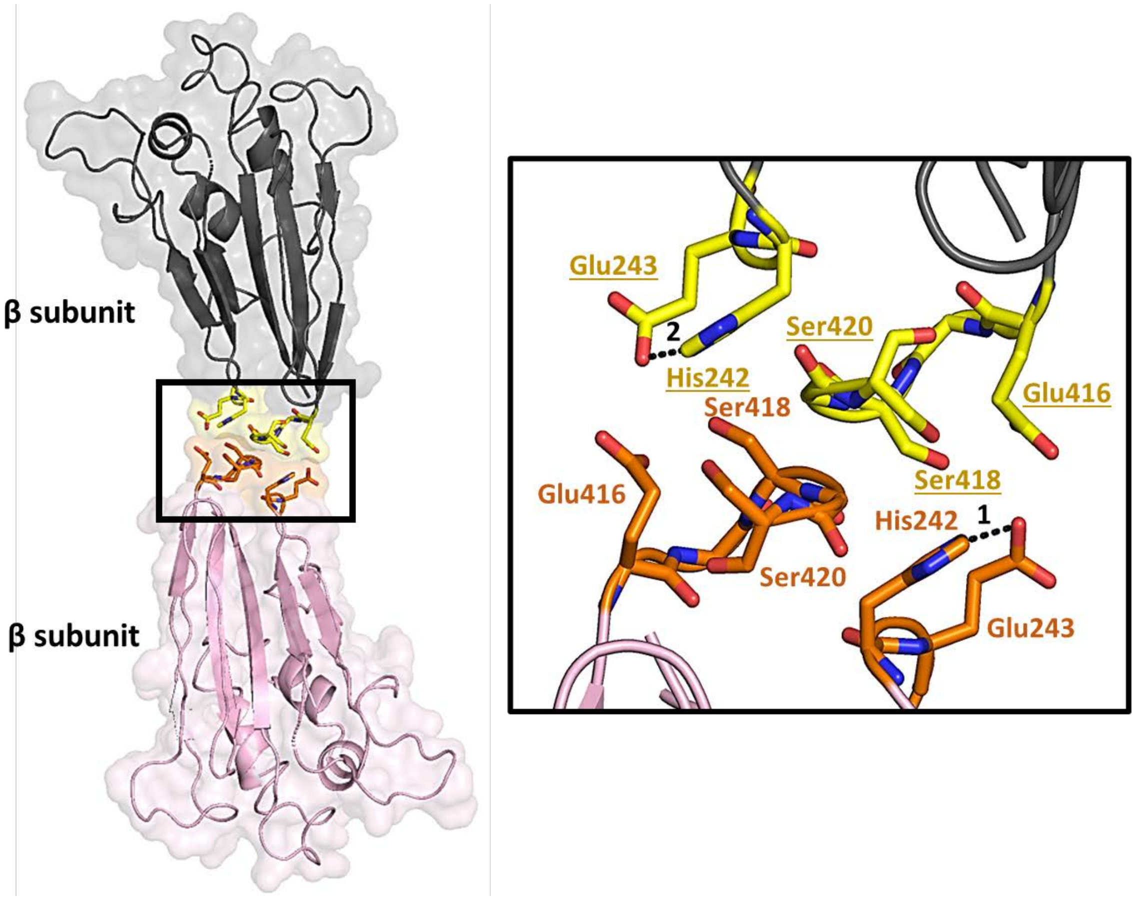


**Figure S23.** Interactions at the linker interface of cpTaspase1$_{\alpha 41\text{-}233/\beta}$. Overview of the linker interface. Cartoon and surface representation of the β subunits molecules involved in the interface. Molecule from the β subunit at lower ring is highlighted light pink and that from the upper ring is highlighted black. The linker interface is indicated by the black box. The black box on the right shows a closer view of the residues at the interface. All residues involved in the interface are shown as sticks in orange and yellow. Contacts at the interface are shown as black dashed lines and numbered according to that in Table S12. For clarity, residue names and numbers from the symmetry related β subunit are underlined.

### Tables

**Table S1.** Comparison of average R.M.S.D. values* between Cp-Taspase1$_{\alpha 41\text{-}183/\beta}$ and the truncated Taspase1 (PDB 2A8J)(14).

| | **C$_\alpha$ atoms (Å)** | **All atoms (Å)** |
|---|---|---|
| Chain A/*α subunit* | 0.384 | 0.524 |
| Chain A/β *subunit* | 0.558 | 0.684 |
| Chain B/*α subunit* | 0.271 | 0.500 |
| Chain B/β *subunit* | 0.558 | 0.684 |

**For consistency, the long helix domain (residues 183-233) were not consider in the analysis.*

**Table S2.** Residues modeled in each of the proteins presented in this study and their comparison to the truncated Taspase1(14).

| **Protein** | **Chain A*** | | **Chain B*** | |
|---|---|---|---|---|
| | ***α subunit*** | ***β subunit*** | ***α subunit*** | ***β subunit*** |
| Activated Taspase1 | 41-155,156-159,160-183,184-233 | 234-349,350-363,364-416,417-420 | 41-156,157-158,159-182,183-233 | 234-351,352-361,362-416,417-420 |
| Cp-Taspase1$_{\alpha 41\text{-}183/\beta}$ | 41-152,153159,160-185,186-233 | 234-350,351-361,362-415,416-420 | 41-152,153158,159-182,183-233 | 234-328,329-337,338-351, 352-361,362-416,417-420 |
| Cp-Taspase1$_{\alpha 41\text{-}233/\beta}$ | 41-228 | 234-351, 352-362, 363-420 | 41-228 | 234-351, 352-362, 363-420 |

**Missing residues in each structure are highlighted red color.*

**Table S3.** Comparison of average R.M.S.D. values* between Cp-Taspase1$_{\alpha 41\text{-}233/\beta}$ and Taspase1 structures.

| | $C_\alpha$ atoms (Å) | All atoms (Å) |
|---|---|---|
| *Cp-Taspase1$_{\alpha 41\text{-}233/\beta}$ vs proenzyme (PDB 2A8I)(14)* | | |
| Chain A/*α subunit* | 0.765 | 1.122 |
| Chain A/β *subunit* | 0.785 | 1.039 |
| Chain B/*α subunit* | 0.662 | 1.084 |
| Chain B/β *subunit* | 0.779 | 1.096 |
| *Cp-Taspase1$_{\alpha 41\text{-}233/\beta}$ vs truncated Taspase1 (PDB 2A8J)(14)* | | |
| Chain A/*α subunit* | 0.646 | 0.981 |
| Chain A/β *subunit* | 0.769 | 1.037 |
| Chain B/*α subunit* | 0.717 | 1.068 |
| Chain B/β *subunit* | 0.720 | 1.092 |
| *Cp-Taspase1$_{\alpha 41\text{-}233/\beta}$ vs Cp-Taspase1$_{\alpha 41\text{-}183/\beta}$* | | |
| Chain A/*α subunit* | 0.614 | 0.900 |
| Chain A/β *subunit* | 0.741 | 0.976 |
| Chain B/*α subunit* | 0.556 | 0.877 |
| Chain B/β *subunit* | 0.932 | 1.189 |

**For consistency, the long helix domain (residues 183-233) were not consider in the analysis.*

**Table S4.** Calculated buried surface area of the catalytic site of Taspase1 structures.

| | **Buried Surface Area (BSA) (Å$^2$)** |
|---|---|
| Cp-Taspase1$_{\alpha 41\text{-}233/\beta}$ chain A | 666.71 |
| Cp-Taspase1$_{\alpha 41\text{-}233/\beta}$ chain B | 754.18 |
| Cp-Taspase1$_{\alpha 41\text{-}183/\beta}$ chain A | 525.65 |
| Cp-Taspase1$_{\alpha 41\text{-}183/\beta}$ chain B | 594.55 |
| Proenzyme (PDB 2A8I) chain A | 519.12 |
| Proenzyme (PDB 2A8I) chain B | 465.52 |
| Truncated Taspase1(PDB 2A8J) chain A | 609.14 |
| Truncated Taspase1(PDB 2A8J) chain B | 623.76 |

**Table S5.** Crystal contacts at the seven interfaces of Cp-Taspase1$_{\alpha 41\text{-}233/\beta}$

| Interface | Interaction[†] Number | Type | Structure 1[#] | Structure 2[#*] | Distance (Å) |
|---|---|---|---|---|---|
| Dimer | 1 | Hydrogen bond | A:LEU125 [O] | B:ARG294 [NH1] | 3.77 |
| | 2 | Hybrid | A:LYS154 [NZ] | B:GLU331 [OE1] | 3.67 |
| | 3 | Hydrogen bond | A:ARG159 [NH2] | B:LEU252 [O] | 3.77 |
| | 4 | Hydrogen bond | A:CYS163 [SG] | B:GLU290 [OE1] | 2.92 |
| | 5 | Hydrogen bond | A:CYS163 [N] | B:GLU290 [OE2] | 3.42 |
| | 6 | Hydrogen bond | A:PHE164 [O] | B:VAL258 [N] | 3.20 |
| | 7 | Hydrogen bond | A:GLU168 [N] | B:GLU168 [OE2] | 3.10 |
| | 8 | Hydrogen bond | A:LEU252 [O] | B:ARG159 [NH2] | 3.28 |
| | 9 | Hydrogen bond | A:ARG257 [NH1] | B:PRO161 [O] | 3.79 |
| | 10 | Hydrogen bond | A:VAL258 [N] | B:PHE164 [O] | 3.14 |
| | 11 | Hydrogen bond | A:TYR264 [OH] | B:ARG294 [O] | 2.46 |
| | 12 | Hydrogen bond | A:GLU290 [OE1] | B:CYS163 [N] | 3.78 |
| | 13 | Hydrogen bond | A:GLU290 [OE1] | B:CYS163 [SG] | 2.64 |
| | 14 | Hydrogen bond | A:GLU290 [OE2] | B:CYS163 [N] | 3.61 |
| | 15 | Hydrogen bond | A:ARG294 [O] | B:TYR264 [OH] | 2.73 |
| | 16 | Hydrogen bond | A:ARG294 [NH1] | B:LEU125 [O] | 3.87 |
| | 17 | Hydrogen bond | A:THR295 [O] | B:ARG299 [NH2] | 2.91 |
| | 18 | Hydrogen bond | A:ARG299 [NH1] | B:THR295 [O] | 2.85 |
| | 19 | Hydrogen bond | A:ARG299 [NH2] | B:GLU300 [E1] | 2.48 |
| | 20 | Salt bridge | A:ARG299 [NE] | B:GLU300 [OE1] | 3.86 |
| | 21 | Salt bridge | A:ARG299 [NH2] | B:GLU300 [OE1] | 2.48 |
| | 22 | Hydrogen bond | A:GLU300 [OE1] | B:ARG299 [NH2] | 2.88 |
| | 23 | Salt bridge | A:GLU300 [OE1] | B:ARG299 [NH1] | 3.44 |
| | 24 | Salt bridge | A:GLU300 [OE1] | B:ARG299 [NH2] | 2.88 |
| | 25 | Salt bridge | A:GLU300 [OE2] | B:ARG299 [NH1] | 3.74 |
| | 26 | Hybrid | A:HIS303 [NE2] | B:GLU300 [OE1] | 3.32 |
| Inter-ring | 1 | Salt bridge | A:GLU313 [OE2] | B:LYS326 [NZ] | 3.91 |

| | | | | | |
|---|---|---|---|---|---|
| | 2 | Hydrogen bond | A:GLU313 [OE2] | B:ASN325 [ND2] | 2.85 |
| | 3 | Hydrogen bond | A:GLU313 [N] | B:GLU321 [OE2] | 3.30 |
| | 4 | Hydrogen bond | A:ALA315 [N] | B:GLN324 [OE1] | 3.69 |
| | 5 | Hydrogen bond | A:HIS316 [O] | B:GLN317 [NE2] | 2.32 |
| | 6 | Hydrogen bond | A:GLN317 [NE2] | B:HIS316 [O] | 2.28 |
| | 7 | Hydrogen bond | A:GLN317 [OE1] | B:LEU320 [N] | 2.51 |
| | 8 | Hydrogen bond | A:LEU320 [N] | B:GLN317 [OE1] | 2.38 |
| | 9 | Hydrogen bond | A:GLN324 [OE2] | B:ALA315 [N] | 3.71 |
| | 10 | Hydrogen bond | A:ASN325 [ND2] | B:GLU313 [O] | 2.36 |
| | 11 | Salt bridge | A:LYS326 [NZ] | B:GLU313 [OE2] | 3.14 |
| | 12 | Hydrogen bond | A:VAL406 [O] | B:GLY411 [N] | 3.66 |
| | 13 | Hydrogen bond | A:GLY411 [N] | B:VAL406 [O] | 3.80 |
| **Histidine** | 1 | Hydrogen bond | A:HIS276 [ND1] | A:GLU308 [OE1] | 2.93 |
| | 2 | Hydrogen bond | A:HIS276 [NE2] | A:GLU308 [OE1] | 3.61 |
| | 3 | Hybrid | B:HIS276 [ND1] | B:GLU308 [OE1] | 2.88 |
| **Linker** | 1 | Hybrid | A:GLU416 [OE2] | A:HIS242 [NE2] | 3.89 |
| | 2 | Hybrid | A:HIS242 [NE2] | A:GLU416 [OE2] | 3.89 |
| **Helix** | 1 | Hydrogen bond | B:HIS53 [NE2] | B:GLU224 [O] | 3.23 |
| | 2 | Hydrogen bond | B:GLU224 [O] | B:HIS53 [NE2] | 3.23 |

**Except for the dimer interface, the structure 2 corresponds to residues from symmetry-related molecules*
*[#]Structures are defined as: chain:residue [atom]*

**Table S6:** Prediction of IDRs in WT Taspase1 sequence.

| Predictor | N-terminal | Middle | C-terminal | Other regions |
|---|---|---|---|---|
| PONDR-FIT | Met1-Ile2 | Thr211-Val235 | Gly411-Ans420 | Lys388 |
| MetaDisorder | Met1-Arg40 | Asn199-Leu232 | Glu416-Asn420 | Ala353-Lys361 |
| MobiDB | Met1-Lys24, Glu29, Glu31 | Lys217-Leu232 | Glu416-Asn420 | Lys154-Leu155, Glu354-Asp356 |
| SPOT-Disorder2 | Met1-Lys39 | Glu207-Ser229 | Ser417-Asn420 | Ser352-Lys361 |
| MoRFpred | | Thr211, Phe213-Leu216 | Glu409, Val412-Arg414, Pro418 | |
